# Multiple pathways for licensing human replication origins

**DOI:** 10.1101/2024.04.10.588796

**Authors:** Ran Yang, Olivia Hunker, Marleigh Wise, Franziska Bleichert

## Abstract

The loading of replicative helicases constitutes an obligatory step in the assembly of DNA replication machineries. In eukaryotes, the MCM2-7 replicative helicase motor is deposited onto DNA by the origin recognition complex (ORC) and co-loader proteins as a head-to-head MCM double hexamer to license replication origins. Although extensively studied in the budding yeast model system, the mechanisms of origin licensing in higher eukaryotes remain poorly defined. Here, we use biochemical reconstitution and electron microscopy (EM) to reconstruct the human MCM loading pathway. Unexpectedly, we find that, unlike in yeast, ORC’s Orc6 subunit is not essential for human MCM loading but can enhance loading efficiency. EM analyses identify several intermediates *en route* to MCM double hexamer formation in the presence and absence of Orc6, including an abundant DNA-loaded, closed-ring single MCM hexamer intermediate that can mature into a head-to-head double hexamer through different pathways. In an Orc6-facilitated pathway, ORC and a second MCM2-7 hexamer are recruited to the dimerization interface of the first hexamer through an MCM-ORC intermediate that is architecturally distinct from an analogous intermediate in yeast. In an alternative, Orc6-independent pathway, MCM double hexamer formation proceeds through dimerization of two independently loaded single MCM2-7 hexamers, promoted by a propensity of human MCM2-7 hexamers to dimerize without the help of other loading factors. This redundancy in human MCM loading pathways likely provides resilience against replication stress under cellular conditions by ensuring that enough origins are licensed for efficient DNA replication. Additionally, the biochemical reconstitution of human origin licensing paves the way to address many outstanding questions regarding DNA replication initiation and replication-coupled events in higher eukaryotes in the future.

## Introduction

Accurate and timely bidirectional replication of genomic DNA is essential for the coordination of genome and cell duplication and the flow of genetic information to progeny. In eukaryotes, this outcome requires that the molecular machinery that replicates DNA (i.e., replisomes) is assembled at hundreds (in budding yeast) to tens of thousands (in humans) origins of replication^1,2^. To accomplish this challenging task and to ensure once-per-cell-cycle genome replication, eukaryotic cells have evolved a cell cycle-regulated, two-step replication initiation process (reviewed in references 3,4). First, origins are licensed by loading two copies of the hexameric <u>M</u>ini<u>C</u>hromosome <u>M</u>aintenance 2-7 (MCM2-7) replicative helicase motor onto replication origin DNA, which results in the formation of stable head-to-head MCM double hexamers^5–7^. In the second initiation step, origin firing, MCM double hexamers mature into active, bidirectional replication forks. Since aberrant origin licensing (both under-licensing and re-licensing of origins) poses a major threat to genome integrity by promoting chromosome instability and tumorigenesis^8^, it is imperative to understand the molecular events needed for successful replisome formation.

Bidirectionality of eukaryotic DNA replication is established during MCM double hexamer formation and relies on the loading of two MCM2-7 hexamers in opposing orientations for dimerization. Biochemical reconstitution of origin licensing with purified *Saccharomyces cerevisiae* (*Sc*) proteins, in conjunction with single-molecule and cryo-electron microscopy (cryo-EM) studies, have illuminated the mechanisms of budding yeast MCM double hexamer assembly (reviewed in references 9-11). In this model (**Fig. 3a**), the heterohexameric <u>O</u>rigin <u>R</u>ecognition <u>C</u>omplex (ORC) binds to specific origin DNA sequences in an ATP-dependent manner^12,13^ and allows association of the co-loader Cdc6^14^. The ORC•Cdc6•DNA complex recruits the first MCM hexamer, along with Cdt1, leading to the formation of an OCCM (ORC•Cdc6•Cdt1•MCM) intermediate, in which the first MCM hexamer topologically encircles DNA^15,16^. Following the release of Cdc6 and Cdt1 – the latter event being coupled to ATP hydrolysis by MCM^17–19^ – the MO (MCM•ORC) intermediate is formed^20^; either the same ORC flips to an inverted secondary DNA binding site at the opposite face of the first loaded Mcm2-7 hexamer^21^, or a second ORC diffuses in to bind at this site^20^. Orc6 plays a critical role in establishing the *Sc*MO – Orc6 stabilizes the first loaded MCM hexamer on DNA and bridges the MCM-ORC interface^20–22^. Further, in the ORC flip model, a long flexible linker in Orc6 likely tethers ORC to MCM while it is repositioned^21^. The MO complex then recruits and loads the second MCM hexamer through the OCCM mechanism^20,21^.

Considerably less is known about how origin licensing occurs in other eukaryotes. While it has been largely inferred that the major steps of the budding yeast MCM loading model are conserved across eukaryotes, there has been little direct evidence to prove that other systems, including human cells, follow the same paradigm. Several significant differences between yeast and human ORC suggest MCM loading may diverge from budding yeast in higher eukaryotes: (1) ORC in most eukaryotes does not bind to a defined origin sequence as in budding yeast^23,24^; (2) in contrast to *Sc*Orc6, metazoan Orc6 proteins (*Drosophila* being an exception) do not stably associate with the core Orc1-5 subunits^25–28^, and (3) they lack the very long flexible linker that is likely utilized by *Sc*Orc6 for ORC flipping and MO formation^27^. Moreover, MCM double hexamers purified from human cells encircle partially melted rather than fully base paired DNA that is seen in the budding yeast counterpart^29–31^, suggesting the possibility of distinct mechanisms for double hexamer formation. Lastly, the lack of full biochemical reconstitution of human replication initiation analogous to budding yeast has hampered progress towards understanding human origin licensing and firing mechanisms. To resolve these conundrums, we developed an *in vitro* reconstituted human origin licensing system with purified proteins and used biochemical experimentation and electron microscopy (EM) to establish how the human MCM2-7 double hexamer is loaded onto DNA.

### *In vitro* reconstitution of human origin licensing

To examine how human MCM2-7 form double hexamers, we developed an *in vitro* reconstitution strategy of human MCM loading similar to that established for budding yeast and *Drosophila* models^5,6,32^. Unlike MCM loading intermediates, MCM double hexamers are salt-stable; thus, MCM loading (i.e., MCM double hexamer formation) can be distinguished from MCM recruitment to DNA (i.e., the assembly of salt-labile MCM loading intermediates) by high- and low-salt washes (**Fig. 1a**). We combined purified human ORC^1–5^, Orc6, Cdc6, Cdt1, and MCM2-7 with bead-immobilized DNA in the absence or presence of nucleotide and assessed MCM recruitment and loading after low-salt and high-salt washes, respectively (**Fig. 1a**). As in budding yeast initiation^5,6^, recruitment of human MCM2-7 and other loading factors was observed with both ATP and the slowly-hydrolysable nucleotide analog ATPγS, while substantial MCM loading was only seen in reactions with ATP, indicating a requirement of ATP hydrolysis for efficient MCM loading (**Fig. 1b**; henceforth, we use only ATPγS for recruitment reactions). MCM2-7 eluted from beads in ATP reactions resembled double hexamers in EM images, confirming successful MCM loading (**Fig. 1c, Extended Data Fig. 1a**).

**Figure 1.**
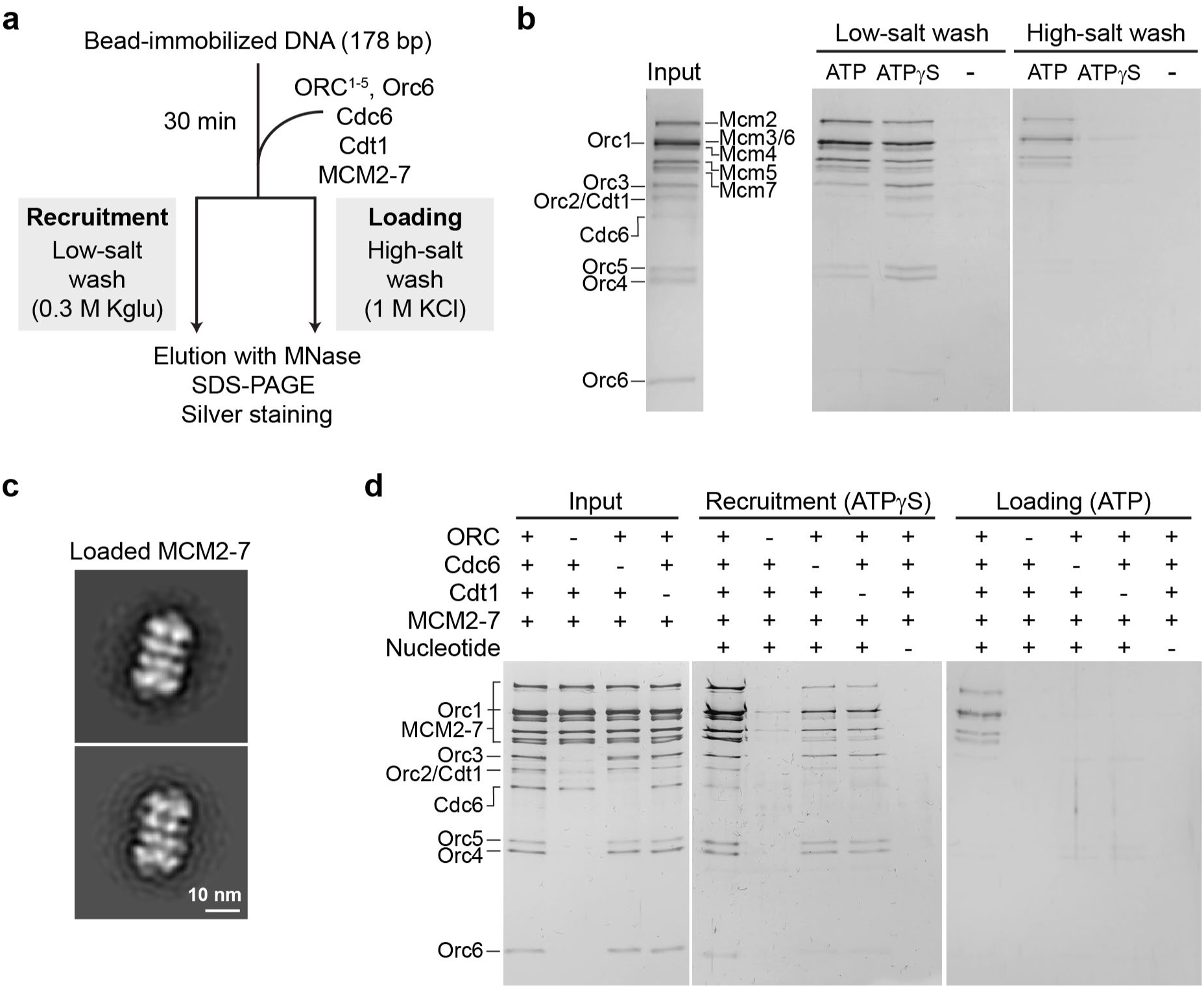
*In vitro* reconstitution of human MCM loading. **a**) Schematic outline of bead-based human MCM recruitment and loading assays. **b**) Nucleotide-dependent recruitment and loading of human MCM2-7. Silver-stained SDS-PAGE gels are shown. Note that low-salt washed reactions with ATP represent a mixture of loading and recruitment, while those with ATPγS reflect mainly recruitment; hence a reduced ORC signal is observed in ATP as compared to ATPγS. **c**) Loaded MCM2-7 complexes are double hexamers. 2D EM class averages of negatively stained particles eluted from beads after high-salt wash. **d**) MCM loading requires ORC, Cdc6, and Cdt1, but some MCM recruitment can occur without Cdc6 and Cdt1. Silver-stained SDS-PAGE gels of inputs and eluates after low-(recruitment) and high-salt wash (loading) are shown. Full-length loading factors were used in these experiments. Kglu – potassium glutamate.

Omission of the initiator ORC or the co-loaders Cdc6 or Cdt1 prevented salt-stable retention of MCM2-7 on DNA; however, a small amount of MCM2-7 could be recruited without Cdc6 or Cdt1, but not without ORC (**Fig. 1d**). While these licensing factors were required for human MCM loading, the intrinsically disordered regions (IDRs) in Orc1, Cdc6, and Cdt1 that promote liquid-liquid phase separation and localization to chromatin^33,34^ were not, confirming that these IDRs are not directly involved in the mechanics of salt-stable double hexamer formation in the human system as previously seen with the fly proteins^32^ (**Extended Data Fig. 1b**). Importantly, addition of human Geminin, a metazoan-specific origin licensing inhibitor that acts by sequestering Cdt1^35,36^, impedes both MCM recruitment and loading, recapitulating a key regulatory aspect of metazoan origin licensing (**Extended Data Figs. 1c-d**). Based on these findings, we conclude that our biochemical *in vitro* reconstitution reflects *bona fide* origin licensing.

To improve the versatility of our reconstituted system, we have utilized truncated constructs in subsequent experiments as IDR removal substantially improved the purification yield of human ORC and Cdt1. Moreover, we have developed a fluorescence-based bead assay to allow more precise quantification of MCM recruitment and loading (**Extended Data Figs. 1e-h**). This workflow relies on MCM2-7 containing GFP fused to the C-terminus of Mcm2, which supported MCM recruitment and double hexamer formation to the same extent as the untagged complex; fluorescence intensity measurements report on the amount of MCM on DNA in recruitment and loading reactions. With these new tools, we set out to dissect how MCM is loaded onto DNA during human origin licensing.

### Orc6 is not essential for but stimulates human MCM loading

Next, we assessed the contributions of individual ORC subunits to human MCM loading. Although ORC is required for MCM loading in yeast, the strict requirement of some ORC subunits for human origin licensing has been debated. Prior studies have reported that certain cancer cell lines can initiate DNA replication at least to some extent without individual initiator subunits like Orc1 or Orc6^37–40^. We found that ORC that lacks Orc1 (ORC^2–5^) could not recruit or load MCM; these defects were not rescued by excess Cdc6, which is an Orc1 paralog and has been suggested to partially compensate for the lack of Orc1^37,40^ (**Extended Data Fig. 2a**). By contrast, Orc6 was not essential for the deposition of human MCM2-7 onto DNA, although Orc6 considerably stimulated MCM loading but not recruitment (**Fig. 2a** and **Extended Data Fig. 2b**). Importantly, MCM double hexamers loaded in the absence of Orc6 appeared structurally similar in 2D class averages to those loaded with Orc6 (**Fig. 2b**). Quantification of the number of salt-stable double hexamers in negative-stain EM images of loading reaction eluates revealed a ∼10-fold increase when Orc6 was included (**Fig. 2c**). The Orc6-mediated enhancement of MCM loading required all Orc6 protein domains *in cis* on the same polypeptide (**Extended Data Figs. 2c-g**). However, the need for Orc6 for efficient salt-stable MCM loading and double hexamer formation could be overcome by increasing concentrations of ORC and Cdc6. Titrating both loading factors in recruitment and loading assays attenuated the differences in DNA-retained MCM-GFP fluorescence and in MCM double hexamer counts in EM images after high-salt washes with and without Orc6 (compared to low ORC/Cdc6 concentrations; **Figs. 2d-f**).

**Figure 2.**
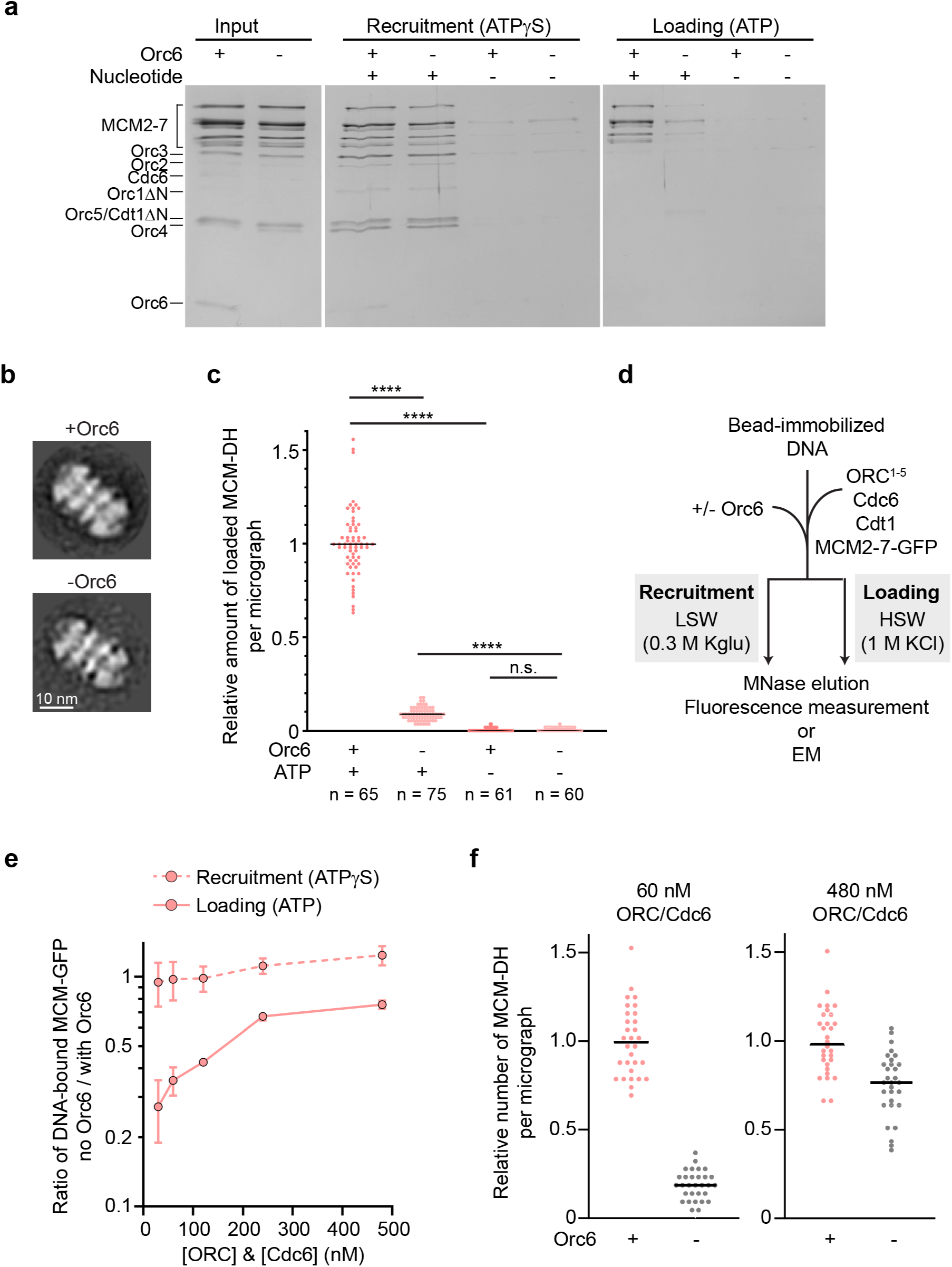
Orc6 is dispensable for human MCM loading *in vitro* but increases loading efficiency. **a**) Silver-stained SDS-PAGE gels of elutions from bead-based MCM recruitment and loading assays with and without Orc6. We note that several ORC subunits (Orc1ΔN, Orc2, Orc6) and Cdc6 stain weakly with silver nitrate. **b**) MCM2-7 complexes loaded onto DNA in the absence of Orc6 are double hexamers and structurally resemble those loaded in the presence of Orc6. 2D EM class averages of negatively stained particles from elutions of loading reactions are shown. **c**) Quantification of MCM double hexamers observed per micrograph from loading reactions with and without Orc6. Micrographs were collected from three independent experiments. The number of counted double hexamers was normalized to the average number of double hexamers per micrograph loaded in the presence of ATP and Orc6. The total number of micrographs (n) analyzed is listed. Statistical significance was calculated using two-way ANOVA with Tukey’s multiple comparisons test (**** p<0.0001, n.s. – not significant). MCM-DH – MCM double hexamer. **d** to **f**) Increasing ORC/Cdc6 concentrations overcomes the requirement for Orc6 in efficient MCM loading. **d**) Schematic of experimental workflow. LSW – low-salt wash; HSW – high-salt wash. **e**) Ratios of GFP fluorescence intensities in elutions of bead-based recruitment and loading reactions without and with Orc6 at a given ORC/Cdc6 concentration. Orc6 was equimolar to ORC^1–5^ when included. Means and standard deviations from three independent experiments are plotted. **f**) MCM double hexamer counts per electron micrograph in elutions from loading reactions at 60 nM and 480 nM ORC/Cdc6 (+/-Orc6) normalized to the respective +Orc6 reaction. Ten micrographs for each of the three independent experiments (total n=30) were analyzed for each condition.

Collectively, these results define the minimal components of our *in vitro* origin licensing system and establish that Orc1, but not Orc6, is essential for human MCM loading. Although Orc6 stimulates this event (especially at low ORC/Cdc6 concentrations), its dispensability was unexpected because *S. cerevisiae* Orc6 is essential for origin licensing^41–43^, raising fundamental questions regarding the loading mechanism of human MCM both in the presence and absence of Orc6.

### Electron microscopy of the human MCM loading pathway

Several intermediates *en route* towards MCM double hexamer formation have been identified in the budding yeast system (**Fig. 3a**). These include an OCCM complex, in which the first MCM hexamer has been deposited onto DNA by ORC, Cdc6, and Cdt1^15,16^. This intermediate transitions into the MO complex, which is stabilized by Orc6 and which recruits the second MCM hexamer^20,21^. To define whether the same intermediates are formed during human MCM loading, we initially analyzed our MCM loading reactions by negative-stain EM. We mixed all essential human loading factors, DNA with ends blocked by streptavidin, and nucleotide, and allowed MCM loading to proceed for 30 minutes before depositing the reaction onto EM grids (**Fig. 3b**). Orc6 was included in these reactions since it stimulates loading (**Fig. 2**). ORC and Cdc6 concentrations (120 nM) were chosen so that they align with nuclear ORC concentrations (∼15-200 nM) estimated by mass spectrometry^44^ and so that loading reactions could be imaged without diluting samples.

**Figure 3.**
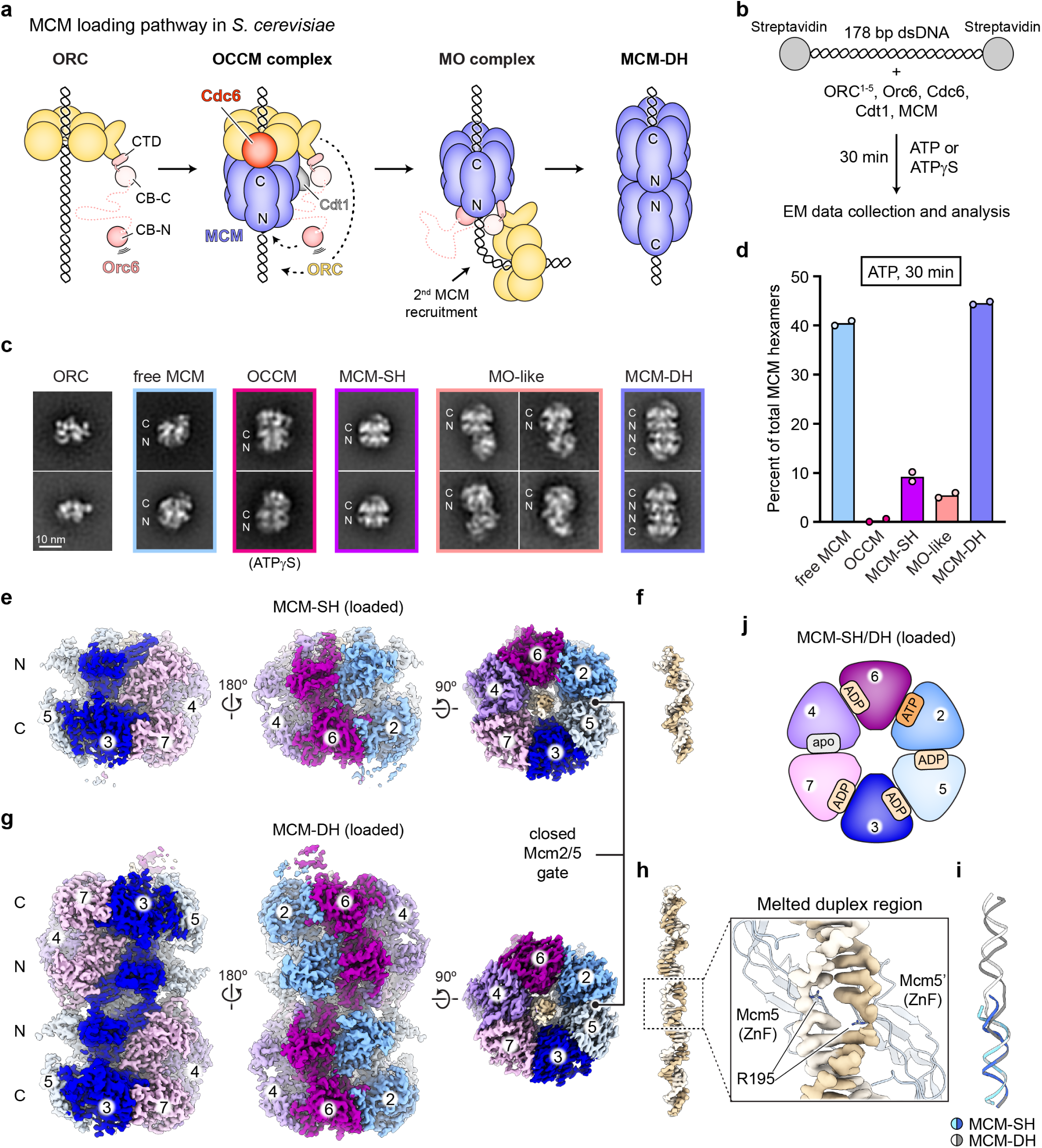
Electron microscopy reveals multiple intermediates during human MCM loading. **a**) Schematic of MCM loading pathway established in *S. cerevisiae*. **b**) Experimental workflow for analyzing human MCM loading reactions by negative-stain EM. **c** and **d**) Representative class averages (in **c**) and quantification (in **d**; datasets from two independent loading reactions) of human MCM2-7-containing assemblies after 30 min loading with ATP. The location of N- and C-terminal tiers in MCM2-7 are marked with N and C. **e** to **j**) Cryo-EM structures of DNA-loaded human MCM single and double hexamers. **e**) Sharpened cryo-EM map of loaded MCM single hexamers (MCM-SH). **f**) Cryo-EM map density for DNA in the central pore of loaded MCM single hexamers. Both DNA strands are fully base paired. **g**) Sharpened cryo-EM map of loaded MCM double hexamers (MCM-DH). **h**) Cryo-EM map density for DNA inside loaded double hexamers. R195 in Mcm5’s zinc finger (ZnF) stabilizes the melting of one base pair at the dimerization interface of *in vitro* loaded MCM2-7. **i**) Superposition of DNA duplexes from loaded MCM single and double hexamers illustrates underwinding and melting of the DNA duplex in MCM-DH. **j**) Nucleotide states in loaded single and double hexamers (viewed from the C-terminal MCM tier). Identical nucleotide states in MCM-SH and DH indicate ATP is hydrolyzed prior to MCM dimerization.

2D classification revealed that almost 50% of MCM hexamers had been incorporated into double hexamer particles (MCM-DH) in the presence of ATP (**Figs. 3c-d**). In addition to free ORC, free (non-loaded) MCM2-7, and MCM-DHs, we also identified a very small number of OCCM intermediates (which were enriched in the presence of ATPγS), as well as ∼10% MCM single hexamer particles (MCM-SH) that closely resembled the hexamers in MCM-DH (**Figs. 3c-d**). MCM-SH class averages were not observed with MCM2-7 alone or at the beginning of the loading reaction, indicating these particles corresponded to loaded single MCM hexamers (**Extended Data Fig. 3a**); we note, however, that the percentage of MCM-SH is most certainly an underestimation since non-side views could not be unambiguously distinguished from free MCM and were therefore counted as free MCM. Strikingly, a subset of class averages was comprised of ORC that was attached to the N-terminal face of an MCM2-7 hexamer (**Figs. 3c-d**). While this organization is reminiscent of the yeast MO intermediate, comparison of these class averages with 2D projections of the *S. cerevisiae* MO suggests the human complex may be structurally distinct, with ORC being oriented differently towards MCM (hence ‘MO-like’; compare **Fig. 3c** and **Extended Data Fig. 3b**). While the molecular complexes observed by negative-stain EM revealed compelling parallels between the yeast and human pathways, the data also suggested differences, particularly with respect to the MO intermediate, that prompted further investigation.

To gain higher resolution information on structural transitions during human MCM loading, we subjected *in vitro* ATP-loading reactions (with Orc6) to cryo-electron microscopy (cryo-EM) analysis. Although the MO-like complexes were of insufficient abundance for cryo-EM structure determination, our data yielded structures of loaded MCM single and double hexamers at 3.4 and 2.8 Å resolution, respectively (**Figs. 3e-h, Extended Data Figs. 4** and **5, Extended Data Table 1**). Several features of these structures are noteworthy. First, the Mcm2/5 gates that serve as the entry site for DNA during loading^45^ are fully closed in both structures, and no additional density is seen near this gate in the MCM-SH cryo-EM map (**Figs. 3e** and **g**); in the *Sc*MO complex, Orc6 latches across the Mcm2/5 gate and stabilizes the loaded MCM single hexamer on DNA^20–22^. This observation suggests that, contrary to yeast, stable Mcm2/5 gate closure may not require Orc6 or formation of the MO complex. Second, DNA is bound in the central pore of both single and double MCM hexamer structures (**Figs. 3f, h-i, Extended Data Fig. 5f**). DNA in the MCM-SH cryo-EM map is fully base paired, whereas one base pair is broken at the hexamer interface in MCM-DH as in the previous open complex structure of endogenous MCM double hexamers from human cells^29^. Thus, DNA underwinding and base pair melting occur during MCM double hexamer formation, and the distinct DNA conformations seen in prior human and budding yeast double hexamer structures are not caused by disparate sample preparation strategies (*Sc*: *in vitro* reconstituted, no DNA melting^30,31^; human: purified from cells, DNA melting^29^) but represent species-specific differences. Third, the ATPase sites in our MCM-SH and MCM-DH structures have identical nucleotide occupancies with five of them having hydrolyzed or released nucleotide (**Fig. 3j**). These postcatalytic states are consistent with ATP hydrolysis by MCM coinciding with single hexamer loading events as established for *S. cerevisiae*^17–19^ but differ from the ATPase configuration in the endogenously purified human MCM-DH^29^, implying that nucleotide can exchange at some of these sites after loading (**Extended Data Figs. 6a-c**).

### Orc6 promotes assembly of an MO-like intermediate during human MCM loading

Our initial negative-stain EM analysis of human MCM loading reactions indicated the formation of an MO-like complex that appeared structurally distinct from the *S. cerevisiae* MO intermediate responsible for recruiting the second MCM hexamer to budding yeast origins (**Fig. 3c, Extended Data Fig. 3b**). Since we were unable to obtain a 3D cryo-EM reconstruction of the human MO-like complex, we used particles in our negative-stain MO-like classes to obtain a low-resolution 3D map of this complex. Docking of ORC and MCM2-7 structures into this map confirmed the different orientations of ORC and MCM as compared to the *Sc*MO (**Extended Data Figs. 3c-e**).

To better understand the role of this distinct, human MO-like complex in origin licensing, we first examined its temporal relationship to other MCM loading assemblies (**Fig. 4a**). Negative-stain EM time course experiments showed that MO-like particles accumulate early during the MCM loading reaction before MCM double hexamers are observed, peak 5 minutes after ATP addition, and then diminish (**Fig. 4b**). MCM-DHs, on the other hand, continuously formed throughout the reaction. The appearance of MO-like particles closely correlated with that of loaded MCM-SH, implying that MO-like complex formation is linked to the loading of single hexamers. Very few OCCM particles were seen, which is consistent with it being a short-lived intermediate as in budding yeast^15,46,47^.

**Figure 4.**
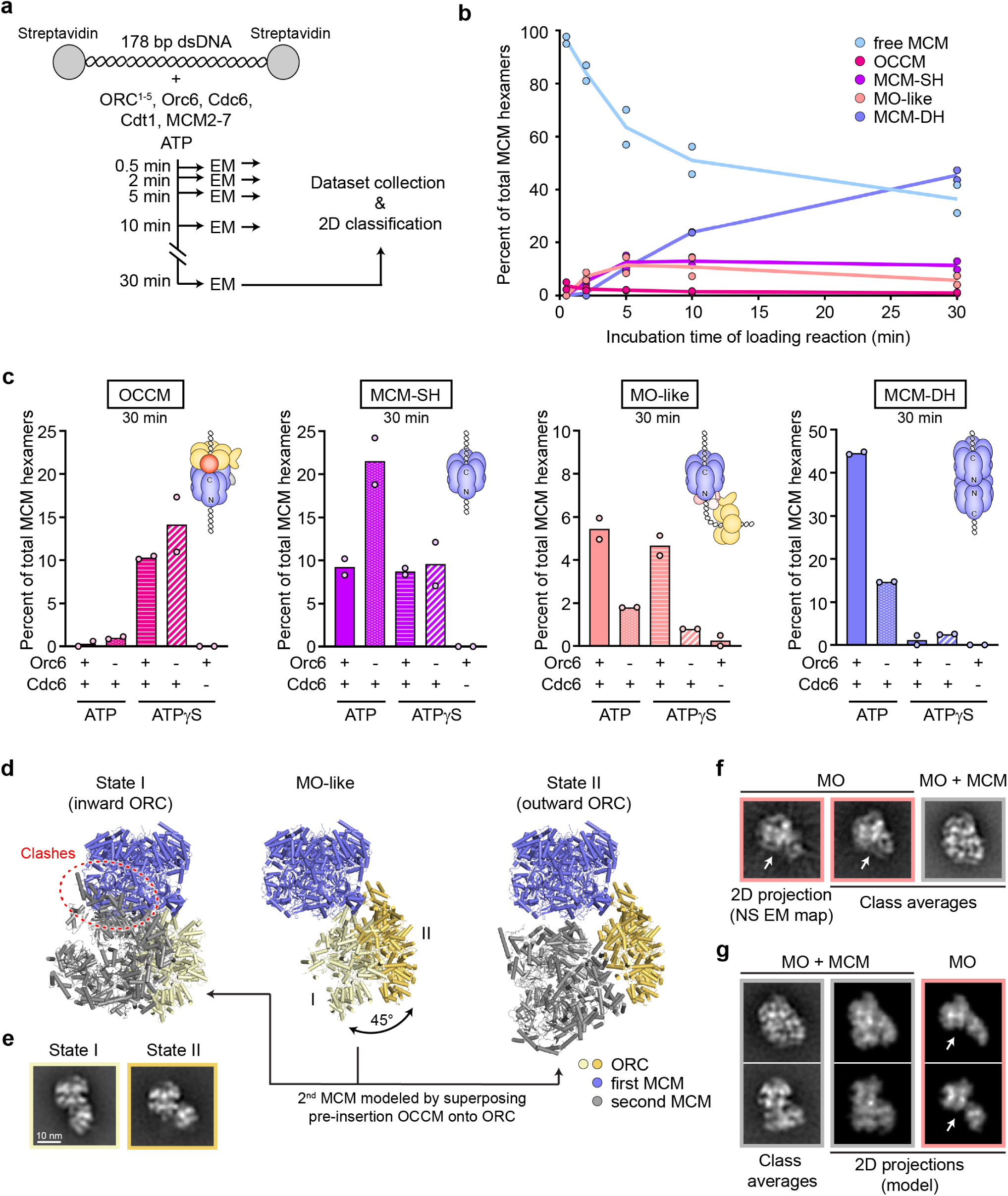
Human Orc6 stimulates assembly of an MO-like complex during MCM loading. **a** and **b**) The MO-like complex is an early MCM loading intermediate. Experimental workflow (in **a**) and quantification of MCM-containing assemblies by negative-stain EM and 2D classification (in **b**) at different time points of loading reactions with ATP. **c**) Quantification (by negative-stain EM) of MCM-containing assemblies after 30 minute-loading reactions with or without Orc6 in ATP or ATPγS. The MO cartoon depicts the organization of the respective yeast complex as in Fig. 3a. In **b** and **c**, negative-stain EM datasets were collected and analyzed from two independent experiments for each time point and condition, respectively. We note that 120 nM ORC/Cdc6 were used in loading reactions for negative-stain EM compared to 60 nM in bead-based loading assays; this modification explains the diminished relative difference in MCM double hexamer counts between -Orc6/+Orc6 conditions here as compared to Fig. 2c (see also **Figs. 2e-f**). **d** and **e**) The human MO-like complex can adopt two distinct conformational states. **d**) Superposition of MCM in both structural models of the human MO-like complex reveals a large swinging motion of ORC (middle; see also **Extended Data Figs. 3c** and **f**). Aligning ORC from the yeast pre-insertion OCCM (PDB 6wgg^48^) onto ORC in the human MO-like complexes indicates that MO state II is compatible with recruitment of a second MCM hexamer (right) but MO state I is not (left). **e**) 2D negative-stain EM class averages for MO-like states I and II. **f** and **g**) A small number of 2D class averages (negative stain) show additional density consistent with a second MCM hexamer being recruited to the human MO. **f**) 2D projection of MO (state II) negative-stain (NS) EM map and two classes corresponding to the same view of the MO with extra density in the right one (MO+MCM). **g)** Comparison of class averages of a putative MO + 2^nd^ MCM complex with 2D projections generated from a modeled complex as in **d** (the model was converted to an EM volume and low-pass filtered to 25 Å before projecting into 2D). Corresponding 2D projections generated from the MO only (state II) model are shown on the right. Since our experimental averages indicated a more compact structure than that obtained when the pre-insertion OCCM is used for modeling, the projections were generated from a model obtained in an analogous manner with the canonical OCCM (PDB 5v8f^16^). We note that some clashes were observed in this model that are likely due to minor inaccuracies in modeling. White arrows in **f** and **g** mark the region in MO class averages or projections that are occupied by a second MCM hexamer in the MO+MCM images.

To establish the function of the human MO-like complex during MCM double hexamer formation more firmly, we next asked whether this complex is involved in a) loading of one or both MCM hexamers onto DNA and b) the Orc6-mediated stimulation of MCM double hexamer formation observed in our system (**Fig. 2**). To this end, we collected negative-stain EM datasets of MCM loading reactions (at 30 min) performed with ATP or ATPγS, and with or without Orc6, and quantified the number of MCM hexamers in OCCM, MCM-SH, MO-like, and MCM-DH 2D classes (**Fig. 4c**). In agreement with bead-based MCM loading assays, reactions without Orc6 supported the assembly of double hexamers with ATP, but at reduced levels. MCM-DH formation was also inefficient in ATPγS, stalling at the OCCM stage as evinced by the strong increase in OCCM particles. Importantly, accumulation of OCCM was not negatively affected in no-Orc6 reactions, indicating that Orc6 is not required for human OCCM assembly; this result is consistent with the efficient recruitment of loading factors to DNA in bead assays in the absence of this ORC subunit (**Fig. 2a, Extended Data** Figs. 2b and f).

By contrast, Orc6 omission led to a ∼2-fold increase in loaded MCM-SH particles in the ATP condition and a strong reduction in the number of MO-like complexes, indicating that the MO-like assembly acts after single hexamer loading onto DNA but prior to double hexamer formation. Moreover, the appearance of MCM-SH and MO-like classes was dependent on OCCM formation and mostly negligible in ATPγS reactions without Cdc6 (an essential component of the OCCM). Surprisingly, substituting ATPγS for ATP in the presence of Cdc6 did not completely block MCM loading at the OCCM stage but led to robust accumulation of MCM-SH and MO-like particles, as well as to some MCM double hexamers. Although we cannot rule out that ATPγS is inefficiently hydrolyzed at some of the MCM ATPase sites, the relatively high number of MCM-SH and MO-like particles suggests that transitioning past the OCCM stage can to some extent be uncoupled from ATP hydrolysis in the human system; however, both events likely coincide in the presence of ATP considering that our DNA-loaded, human MCM-SH cryo-EM structure is in the postcatalytic state (**Fig. 3j; Extended Data** Fig. 6c). ATP hydrolysis, on the other hand, is required for efficient maturation of single into double MCM hexamers, potentially by inducing a conformational change needed for stable dimerization. Collectively, these results argue that the MO-like complex is a *bona fide* MCM loading intermediate that acts after the OCCM in the loading pathway and whose assembly is facilitated by Orc6.

In *S. cerevisiae*, the MO intermediate coordinates the recruitment of a second MCM hexamer to an already loaded first hexamer in the correct orientation for head-to-head double hexamer formation^20^. Although we initially presumed that the human MO-like complex performed an analogous function, this assumption is challenged by the distinct orientation of ORC with respect to the MCM2-7 hexamer (**Extended Data Figs. 3c-e**); indeed, modelling of a second MCM hexamer by superposition of ORC from the budding yeast pre-insertion OCCM structure^48^ onto the docked ORC model in the negative-stain EM 3D reconstruction of the human MO-like complex revealed major clashes between the two MCM2-7 rings (**Fig. 4d**, state I on the left). Closer inspection of MO-like class averages in our various negative-stain datasets, however, uncovered discernable flexibility of ORC, positioned either near the center or near the edge of the N-terminal face of MCM2-7 (**Fig. 4e, Extended Data Video 1**). Combining MO-like particles from multiple datasets yielded a low-resolution map of a second MO-like state, in which ORC had rotated outwards by ∼45 degrees, opening the space between ORC and MCM that can now accommodate a second hexamer as part of a pre-insertion OCCM (**Fig. 4d**, state II on the right; **Extended Data Figs. 3f-h**). Moreover, 2D projections generated from this modeled MCM-OCCM complex resembled a few averages that could not be accounted for by the other loading intermediates considered so far (**Figs. 4f-g**).

These analyses provide further evidence that the human MO-like complex is an on-pathway loading intermediate and that, despite their distinct architectures (human and *Sc*ORC use completely different regions to bind MCM, **Extended Data Figs. 3d** and **g**), *Sc*MO and the human MO-like complex may serve analogous roles: recruiting the second MCM hexamer to the first loaded MCM and ensuring it is loaded in the correct orientation to afford double hexamer assembly. Given the functional parallels between the budding yeast and human complexes, we will refer to the human one as MO henceforward.

### Human MCM double hexamers can form in an MO-independent manner by dimerization of independently loaded hexamers

Given our findings that human Orc6 enhances MO assembly but is dispensable for MCM double hexamer formation, we considered the possibility of an alternative, MO-independent MCM loading pathway. This rationale was reinforced by our observation that human MCM2-7 has a strong propensity to dimerize during protein purification in the absence of other loading factors as demonstrated by its early elution in size exclusion chromatograms (**Extended Data Fig. 7a**). Dimers can be clearly seen both in negative-stain EM and cryo-EM class averages, and they are structurally distinct from loaded MCM double hexamers (**Extended Data Figs. 7b-c**); we will thus refer to this double hexameric form of MCM2-7 as MCM dimer to distinguish it from the DNA-loaded double hexamer.

We determined the cryo-EM structure of this MCM dimer at a resolution of 3.8 Å, which was improved to 3.5 Å by symmetry expansion and local refinement (**Extended Data Figs. 7d-h** and **8a-e, Extended Data Table 1**); we used chemical crosslinking to reduce flexibility between the two MCM2-7 rings without affecting the overall architecture of the MCM dimer (compare **Extended Data Figs. 7c** and **e**). In this structure, two MCM2-7 hexamers stack onto each other with the N-terminal domains facing (**Fig. 5a**). Both MCM2-7 rings adopt a left-handed lock-washer conformation with each of the Mcm2/5 gates, which serve as the site for DNA entry during MCM loading^45^, clearly open. This configuration restricts the dimerization interface to Mcm3, Mcm5, and Mcm7, which engage the other hexamer using contacts that are similar to those in the loaded MCM double hexamer (**Fig. 5b** and **Extended Data Figs. 8f-i**). We observe clear density for nucleotide at five of the ATPase sites (all ATP except for ADP at the Mcm5-3 site), while the sixth one (Mcm2-5) is empty (**Extended Data Figs. 6a** and **d**). These structural features, in conjunction with the absence of DNA in the central pores, are consistent with MCM2-7 in the dimer being in a pre-loading state and distinct from the MCM double hexamer.

**Figure 5.**
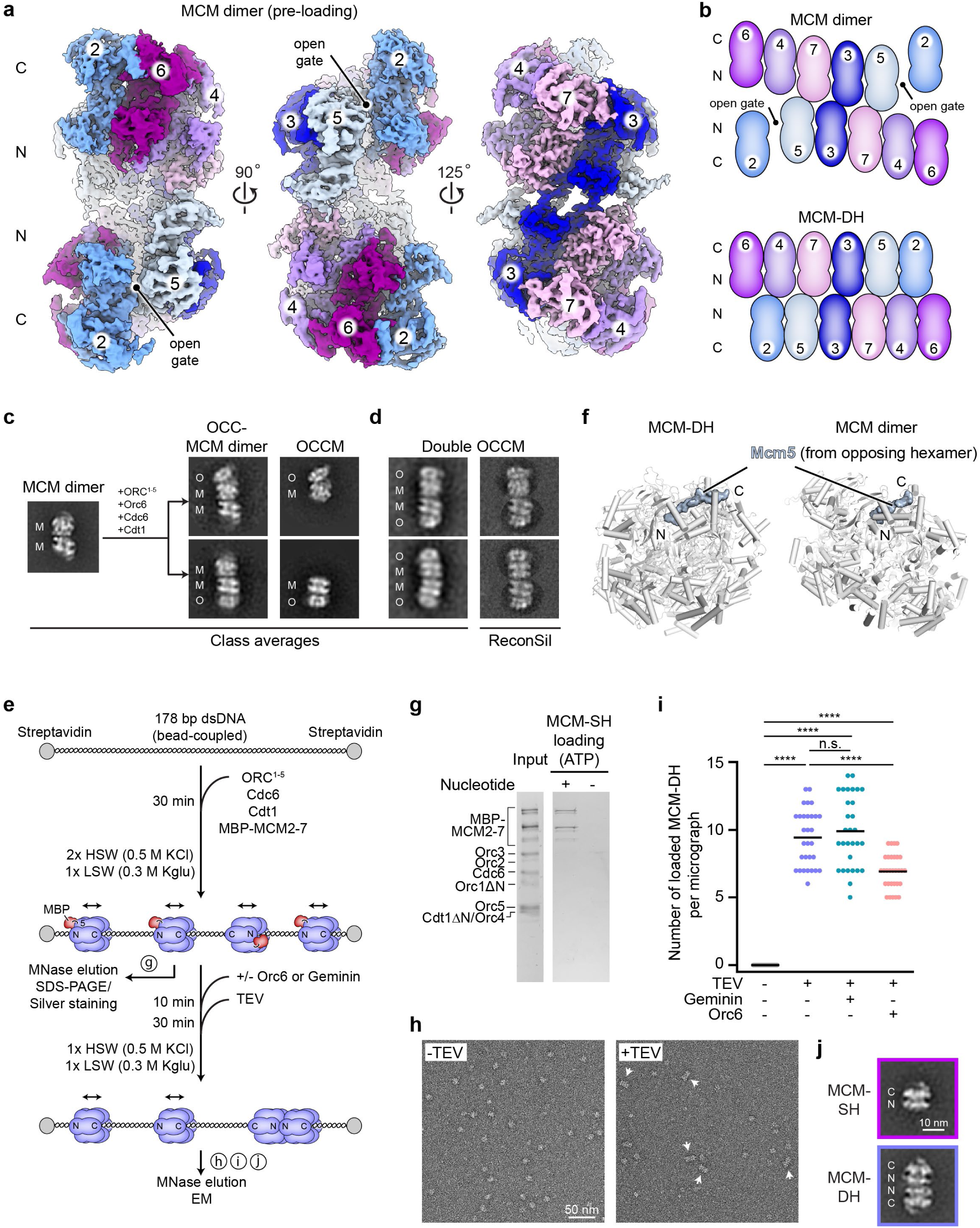
Human MCM2-7 hexamers have a propensity to dimerize without other loading factors and can mature into double hexamers by dimerization of independently loaded hexamers. **a**) Cryo-EM structure of the human MCM2-7 dimer with an open gate in each of the hexamers. **b**) Schematic of subunit interactions in the MCM dimer (pre-loading) and MCM double hexamer (loaded). The MCM dimer is stabilized by contacts between Mcm3, Mcm5, and Mcm7 in both hexamers across the dimerization interface. **c** and **d**) MCM dimers can associate with other loading factors. **c**) Negative-stain EM 2D class averages of a free MCM dimer, ORC bound to one hexamer in MCM dimers (likely forming an ORC•Cdc6•Cdt1 (OCC)•MCM dimer complex), and of the canonical OCCM intermediate. Reactions were performed for 5 minutes with ATPγS without pre-heating MCM2-7. **d**) Class averages of double OCCMs and *in silico* reconstitution (ReconSil^20^) images of double OCCMs generated by superposition of single OCCM class averages from reactions done with ATPγS onto electron micrographs. **e** to **j**) Independently loaded MCM single hexamers can dimerize on DNA to form double hexamers after removal of other loading factors. **e**) Overview of experimental setup. MCM single hexamers are first loaded onto DNA in the absence of Orc6. MCM double hexamer formation is prevented by an N-terminal MBP tag on Mcm5, which is cleaved off with TEV after removal of ORC, Cdc6, Cdt1, and non-loaded MCM2-7. Double hexamer formation is evaluated by negative-stain EM. Letters in circles refer to figure panels with results of respective experiments. **f**) The Mcm5 N-terminus of one MCM hexamer is buried in the adjacent hexamer in both the MCM dimer and loaded double hexamer. N and C mark the respective termini of the Mcm5 fragment. **g**) Silver-stained SDS-PAGE gel of loading factor mix (Input) and loaded MCM single hexamers after removal of other loading factors and before TEV cleavage. **h)** Subregion of electron micrograph (negative stain) of loaded MCM particles treated with or without TEV and eluted with MNase. Double hexamers (white arrowheads) are only observed after TEV treatment. **i**) Quantification of MCM double hexamer numbers per micrograph. 10 micrographs each in a total of three independent experiments were analyzed per sample (total n=30). Black lines represent means. Statistical significance was determined by two-way ANOVA and Tukey’s multiple comparison tests. **** p<0.0001, n.s. – not significant. **j**) 2D class averages of negatively stained MCM double hexamers formed by dimerization of single hexamers after MBP removal by TEV cleavage as in **h** and **i**.

What might be the purpose of MCM dimerization prior to loading? Of the isolated MCM assemblies (i.e., not loaded) structurally characterized to date, the human complex appears to be unique in this regard. *Drosophila, E. cuniculi,* and *S. cerevisiae* hexameric MCM rings adopt similar conformations as the human counterpart but do not dimerize before loading^5,6,49–51^. We reasoned that the strong propensity of human MCM to dimerize might facilitate MCM double hexamer formation through an MO-independent pathway, which could become more important without Orc6 and in the absence of specific DNA sequences at human origins that could direct ORC to DNA in an inverted orientation for second hexamer loading.

There are several possibilities for how MO-independent MCM double hexamer formation could occur: *1) MCM could be recruited and loaded onto DNA as a preformed dimer.* Our data suggest that this event is infrequent under our experimental conditions. While MCM dimerization is not affected by nucleotide or the addition of Cdt1 (which in yeast but not humans binds MCM to form heptamers preceding origin recruitment^5,52,53^, MCM dimers are temperature sensitive, at least at nanomolar protein concentrations (**Extended Data Figs. 9a-b**); at 37°C (the temperature used for loading reactions), the equilibrium shifts towards the single MCM2-7 hexamer, although some dimers remain (**Extended Data Fig. 9b**). Moreover, while we observe 2D class averages in which ORC is bound to one of the hexamers in an MCM dimer (OCC-MCM dimer) in an analogous manner as seen in the OCCM, such particles are rare, and we don’t know whether these complexes can proceed towards double hexamers (**Fig. 5c**). *2) MCM2-7 could dimerize within the context of loading intermediates prior to MO formation, i.e. at the OCCM stage.* Interestingly, we observe OCCM particles that appear to interact through the N-terminal face of MCM to form “double OCCMs”, especially at higher ORC/Cdc6 concentration (**Fig. 5d**). These double OCCMs could arise either by ORC recruitment to OCC-MCM dimers or by DNA sliding and dimerization of two independently formed OCCMs. Since single OCCM classes are much more abundant than OCC-MCM dimers, we favor the latter scenario. *3) MCM double hexamers could form by sliding and dimerization of independently loaded single MCM hexamers, bypassing the MO intermediate but facilitated by Mcm5/3/7 co-association between hexamers*. In contrast to OCC-MCM dimers and double OCCMs, DNA-loaded single MCM hexamers were observed in our cryo-EM data and were frequently seen in negative-stain class averages of loading reactions (**Figs. 3d-e** and **4c**), indicating that these complexes may be sufficiently stable on DNA for encountering a second loaded hexamer through sliding (note that yeast loaded single hexamers can slide on DNA^54–56^). We therefore decided to test this third premise directly.

To determine whether human MCM double hexamer formation could occur by dimerization of independently loaded hexamers without an MO intermediate, we developed a biochemical assay in which single MCM hexamers could be loaded onto bead-coupled DNA but not form double hexamers until after other loading factors had been removed (**Figs. 5e-f**). To prevent MCM dimerization, we added an MBP tag, followed by a TEV protease site, to the N-terminus of Mcm5, which is flexible in the single MCM hexamer but buried within the opposing hexamer in the MCM dimer and loaded double hexamer (**Fig. 5f**). As expected, this complex purified as a single hexamer, was loaded onto DNA exclusively as single hexamers (0.5 M KCl wash), and did not form salt-stable double hexamers (1 M KCl wash; **Extended Data Figs. 7a, 9c-f**). Removal of the large MBP moiety by TEV digestion prior to loading reactions restored salt-stable loading, indicating that the cleaved MCM hexamer is fully functional (**Extended Data Figs. 9d-f**).

We then loaded MBP-tagged single hexamers onto DNA in the absence of Orc6 to mitigate MO formation and washed away free and DNA-bound loading factors other than loaded MCM with 0.5 M KCl buffer (**Figs. 5e** and **g, Extended Data Fig. 9g**). If these independently loaded single hexamers can dimerize, we should observe the appearance of double hexamers in EM images of eluates after TEV addition. Indeed, TEV-treated samples had on average 10 double hexamers per electron micrograph, whereas mock-treated ones had none (**Figs. 5h-j**). Inclusion of Geminin, a Cdt1 inhibitor, during TEV cleavage did not alter the number of MCM-DHs observed, consistent with double hexamers not arising from new MCM loading (**Fig. 5i**). Surprisingly, addition of excess Orc6 after single hexamer loading inhibited double hexamer formation after TEV cleavage (**Fig. 5i**). This result confirmed that double hexamerization is MO-independent in this experiment (if the MO were involved, for example because a small amount of ORC was retained after washes, we would expect to see an increase in MCM-DHs with Orc6), but also suggests that Orc6 can act as a block to dimerization. Indeed, AlphaFold Multimer^57,58^ predicts that human Orc6 binds the N-terminal domain of Mcm6, which could sterically impede dimerization.

Collectively, these results demonstrate that human origin licensing can occur in an MO-independent manner by dimerization of independently loaded MCM2-7 complexes (likely after sliding on DNA), thereby uncoupling the second MCM2-7 loading event from MCM dimerization. This loading pathway can partially explain the rescue of efficient MCM loading in the absence of Orc6 at higher ORC/Cdc6 concentrations (**Figs. 2e-f**) as more MCM single hexamers would be expected to be loaded onto a given DNA, increasing the likelihood of productive MCM encounters.

### Disease-linked mutations in Orc6 impede human MO formation and MCM loading

Our discovery of multiple routes towards human MCM loading and double hexamer formation raises questions pertaining to their individual contributions to origin licensing *in vivo*. Previous work has shown that depletion of Orc6 in U2OS osteosarcoma cells did not impede chromatin association of MCM^39^ (although it is unknown whether these are single or double hexamers), indicating that Orc6 (and by inference high levels of the MO intermediate) may not be essential for efficient origin licensing in humans. Nonetheless, mutations in Orc6 and other licensing factors, are associated with a form of primordial dwarfism, Meier-Gorlin syndrome (MGS), that is caused by an impediment of DNA replication^59^. We thus asked whether MGS patient variants of Orc6 have a direct impact on MCM double hexamer formation in our *in vitro* MCM loading system, focusing on missense mutations in Orc6’s CB-N region (K23E^60^) and C-terminal domain (Y232S^61^) (**Fig. 6a, Extended Data Fig. 2d**).

**Figure 6.**
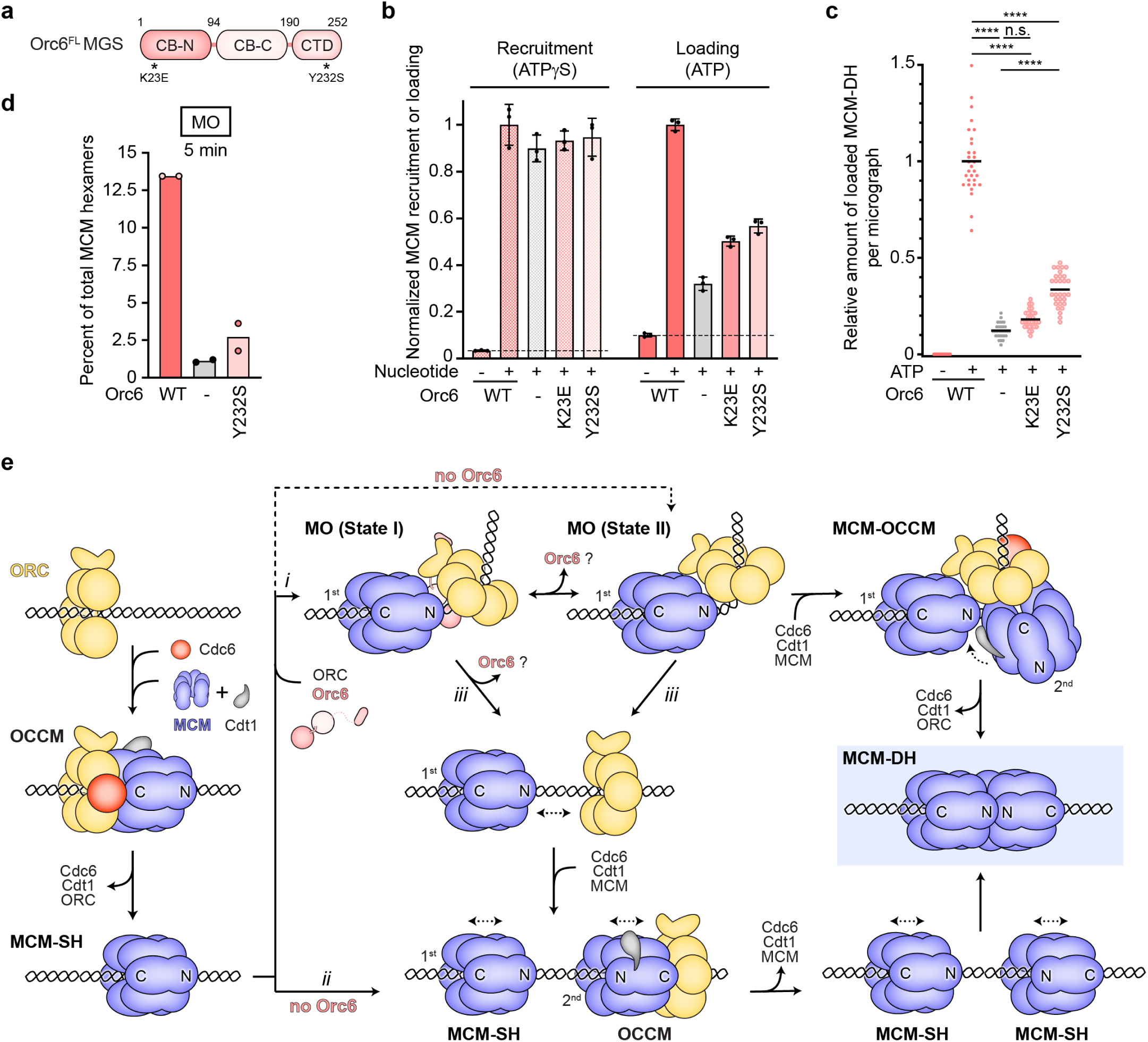
Disease-linked mutations in human Orc6 impair formation of the MO complex and MCM loading. **a**) Domain architecture of human Orc6 illustrating the location of mutations associated with Meier-Gorlin syndrome (MGS). **b** and **c**) MGS mutations in Orc6 (K23E and Y232S) impede efficient salt-stable MCM double hexamer formation but not MCM recruitment. **b**) The means and standard deviations of MCM-GFP fluorescence measured in elutions of three independent recruitment and loading reactions (1 M KCl wash) are plotted. The fluorescence signal was normalized to the average fluorescence signal obtained with wildtype (WT) Orc6. **c**) Quantification of MCM double hexamers observed in electron micrographs (negative stain) of loading reaction elutions (n=30 with 10 micrographs per each of three independent repeats). Statistical significance was determined with two-way ANOVA and Tukey’s multiple comparisons tests. **** p<0.0001, n.s. – not significant. We note that the fluorescent assay in **b** slightly overestimates MCM loading efficiency (without Orc6 or with Orc6 mutants) since loaded single hexamers also contribute to the GFP signal. **d**) The Orc6^Y232S^ MGS mutation impairs MO complex formation. MO particles were quantified by negative stain-EM and 2D classification of MCM loading reactions. A 5-minute reaction time was chosen because the MO intermediate is the most abundant at this time point (see Fig. 4b). Datasets were collected from two independent loading reactions. **e**) Model for human origin licensing pathways. Orc6 is not depicted during OCCM assembly since it is not required for this MCM loading step and only weakly associates with human ORC. We note that pathway *ii* could also occur by sliding of two OCCMs. Dashed line signifies inefficient MO formation without Orc6 (likely MO state II since we do not observe 2D classes characteristic of state I in this condition; **Extended Data Fig. 3i**). Dotted arrows indicate sliding.

Our results demonstrate that both Orc6-MGS mutants display an MCM loading defect in bead-based loading assays and significantly compromise formation of salt-stable double hexamers (**Figs. 6b-c**). Moreover, analyses of intermediates in MCM loading reactions in the presence of ATP by negative-stain EM showed a strong reduction in MO complex formation for Orc6-Y232S (**Fig. 6d**). These findings reinforce the conclusion that Orc6 is not strictly essential for human origin licensing but that it increases MCM loading capacity through an MO-mediated loading pathway that is relevant to growth and cell proliferation at least during certain stages of development.

## Discussion

Bidirectional replication fork establishment is a universal feature of DNA replication initiation across the domains of life but is achieved through a variety of mechanisms^62^. In eukaryotes, this critical event requires two MCM complexes to be deposited onto DNA in opposite orientations as a head-to-head double hexamer during origin licensing. How this specific MCM architecture is established efficiently in eukaryotes, particularly in those other than a small group of *Saccharomycetales* species that rely on specific origin DNA sequences to guide the directional loading of two MCM complexes, has remained a fundamental, unresolved question. In this study, we elucidate the mechanisms underlying MCM loading in the human system, revealing unexpected flexibility in loading strategies that are likely the norm in a wide range of eukaryotes.

Our findings afford the following model summarizing human origin licensing pathways (**Fig. 6e**). The initiator ORC, with the help of Cdc6 and Cdt1, first recruits and loads a single MCM hexamer onto DNA through an OCCM intermediate similarly to budding yeast^15,47^. Human Orc6 does not appear to contribute to this loading event (**Figs. 2** and **4**), a conclusion in agreement with DNA-bound metazoan ORC structures that do not show Orc6 contributing to DNA binding and bending as in *S. cerevisiae*^13,32,48,63^. Our results further argue that single loaded human MCM hexamers can then mature into double hexamers in several different ways. In the presence of Orc6, an MO-dependent pathway is likely preferred (**Fig. 6e*i***), in which Orc6 facilitates ORC binding to the N-terminal face of the first loaded MCM, forming the MO complex (**Fig. 4**). We postulate that ORC is recruited from solution rather than through an ORC flip as seen in budding yeast; this assumption is borne out by human Orc6’s weak affinity for ORC^1–5^ and by the lack of a sufficiently long Orc6 linker for establishing a tether with the N-terminal surface of the MCM ring^25–27^. The resultant human MO can exist in two different conformations (MO states I and II, **Fig. 4**), only one of which (MO state II) is able to accommodate the recruitment of a second MCM hexamer; the other conformation may correspond to a pre-state that relies on Orc6 to first localize ORC^1–5^ to the first MCM hexamer before shifting into a conformation (possibly coupled to the dissociation of Orc6) poised for second MCM recruitment. Without Orc6, MO formation is less efficient, likely due to a low affinity of ORC^1–5^ alone for MCM (**Figs. 2, 4,** and **6**). The outcome of this MO-dependent, Orc6-facilitated pathway is analogous to MO-mediated MCM loading in *S. cerevisiae*, although the architecture of the MO and role of Orc6 have diverged from yeast to humans.

In an alternative, Orc6-independent pathway, human MCM double hexamers can form by dimerization of independently loaded MCM single hexamers (**Fig. 6e*ii***), bypassing the MO intermediate (**Fig. 5**). This loading mechanism likely requires sliding of OCCMs (or OCMs after Cdc6 release) and/or single MCM hexamers on DNA, properties that have been reported for the yeast intermediates^54–56^. We speculate that human MCM double hexamer formation through MCM sliding and dimerization is enhanced by the strong propensity of human MCM2-7 to dimerize through Mcm5/3/7. We note that MCM double hexamer formation through an MO-dependent pathway and through sliding/dimerization are not mutually exclusive; it is also conceivable that ORC might disengage from MCM after MO assembly but prior to loading the second MCM (**Fig. 6e*iii***). In this regard, the MCM-ORC interactions in the MO would serve to recruit and position ORC on DNA in the correct direction for loading the second MCM hexamer, overcoming the need for specific DNA sequences to orient ORC as in *S. cerevisiae*^12,13,20,54^. Such a mechanism would also reconcile the existence of a human MO complex and the strong human MCM dimerization interface.

Are MO-dependent and -independent loading pathways used interchangeably in human cells, and in metazoa more generally, or is one dominant? Metazoa lack sequence specificity in origin recognition^1,2^, suggesting that origin licensing via the MO-independent pathway has a mere 25% chance for productive head-to-head double hexamer formation, assuming deposition of MCMs in random orientations. However, human cells depleted of Orc6 can sustain normal levels of MCM on chromatin (although it is unclear whether these MCMs are single or double hexamers)^39^. Likewise, *Xenopus* Orc6 is not required for efficient origin licensing^28^, and *C. elegans* does not have an Orc6 ortholog. These observations argue for an important role of MCM dimerization after uncoupled single hexamer loading events. Alternatively, it is also possible that chromatin may help in directing the orientation of MCM loading, or that the local concentrations of loading factors near origins are sufficiently high (e.g., due to liquid-liquid phase separation of initiation factors^33,34^) to promote effective Orc6-independent MO assembly (**Fig. 4c** and **Extended Data Fig. 3i**). Nonetheless, Meier-Gorlin syndrome-associated Orc6 mutants are deficient in both MO and double hexamer formation (**Figs. 6a-d**), defects that are likely responsible for the reduced origin licensing capacity in patient cells and lethality in flies^64,65^, reinforcing the importance of Orc6-stimulated MCM loading. These seemingly disparate observations can be reconciled by the flexibility in MCM loading pathways observed biochemically in our reconstituted system and argue that not all origins in metazoan cells are licensed through the exact same mechanism. By contrast, budding yeast likely experiences a strong evolutionary pressure to license most if not all of its origins by an Orc6- and MO-dependent mechanism to ensure efficient origin licensing control by CDK phosphorylation of Orc6 (and Orc2), which inhibits MO assembly and origin re-licensing in S-phase^22,66,67^. The diverged roles of Orc6 in budding yeast and metazoa likely reflect an evolutionary adaptation to both sequence-independent origin specification and a reliance on Orc6- and MO-independent means to inhibit origin re-licensing in S-phase in higher eukaryotes^3,4,68^.

A second unexpected finding of our studies is the ability of human MCM2-7 to dimerize prior to loading (**Figs. 5a-b**). MCM self-association has been reported for endogenous human MCM2-7, but the nature of the complex formed had been unclear^69^. While there is currently no evidence to support that these MCM dimers can be directly loaded onto DNA, we speculate that the Mcm5/3/7 inter-hexamer interface observed in the dimer may represent the initial contact during double hexamer formation. This interpretation is congruent with single-molecule studies in yeast showing that interactions between two MCM hexamers (involving Mcm7) are established before the loading of the second MCM, or Mcm2/5 gate closure, is completed^19,70^. The strong propensity of MCMs to co-associate through Mcm5/3/7 is likely to benefit the dimerization of independently loaded hexamers as well (**Figs. 5e-j**), facilitating alignment of the correct inter-hexamer register.

In summary, our results demonstrate that human cells employ multiple pathways to license origins and provide key insights into how origin licensing mechanisms have diverged in higher eukaryotes. The plasticity of human origin licensing uncovered here is strongly reminiscent of the known flexibility in mammalian origin selection^1^, and likely functions to provide resiliency against various cellular challenges and conditions, including replication stress and development programs. Moreover, this *in vitro* reconstitution system of human origin licensing represents a critical step towards reconstituting the entire replication initiation pathway in higher eukaryotes, enabling studies of MCM activation mechanisms, epigenetic regulation of initiation, and replication-coupled events in the future.

## Materials and Methods

### Protein constructs and baculovirus generation

Full-length and truncated *Homo sapiens* (*Hs*) ORC^1–5^ were reconstituted in insect cells by infection with baculoviruses expressing multiple ORC subunits. To generate ORC-containing baculovirus vectors, full-length ORC subunits or N-terminally truncated Orc1 (amino acids 400-861, referred to as Orc1ΔN) were first cloned into a ligation-independent-cloning (LIC)-compatible pFastBac vector (Macrolab, University of California Berkeley, USA)^71^. For purification, a hexa-histidine (6xHis) tag and a maltose binding protein (MBP) tag, both followed by a tobacco etch virus (TEV) protease cleavage site, were added to the N-termini of Orc1 and Orc4, respectively. For expression of *Hs*ORC^2–5^, the 6xHis-TEV tag was moved to the N-terminus of Orc2. Subsequently, ORC subunits were combined into a pFastBac-derived BioBricks MultiBac vector (Macrolab, University of California Berkeley, USA) by subcloning different combinations of *Hs*Orc1-5 genes^71^. Bacmids were generated in DH10Bac cells and used for transfections of Sf9 cells with Cellfectin II (Thermo Scientific Fisher). Baculoviruses were amplified for two rounds in Sf9 cells to obtain high-titer viruses for infection of Hi5 cells for large-scale expression.

Full-length wildtype (aa 1-252), MGS-mutant (K23E and Y232S) and truncated (CB-N: aa 1-94, ΔCTD: aa 1-190, CB-C: aa 95-190, and ΔCB-N: aa 95-252) *Hs*Orc6 were cloned as N-terminal 6xHis-TEV fusions into a LIC-converted, pET-derived *E. coli* expression vector (Macrolab, University of California Berkeley, USA). The full-length construct was used as a template to generate expression vectors encoding Orc6 MGS-mutants (Orc6^K23E^ and Orc6^Y232S^) by site-directed mutagenesis. Mutations were verified by DNA sequencing.

*Hs*Cdc6 was expressed in insect cells. The coding sequences of full-length or N-terminally truncated Cdc6 (Cdc6ΔN, amino acids 134-560) were cloned into the LIC-compatible pFastBac vector (Macrolab, University of California Berkeley, USA) with an N-terminal 6xHis-MBP tag and a TEV cleavage site. Bacmid generation and baculovirus amplification were performed as described for *Hs*ORC^1–5^.

Full-length human Cdt1 was expressed in insect cells, while truncated *Hs*Cdt1 was expressed in *E. coli*. The coding sequence of full-length Cdt1 was cloned into the LIC-compatible pFastBac vector (Macrolab, University of California Berkeley, USA) with an N-terminal 6xHis-MBP tag and a TEV cleavage site. Bacmid generation and baculovirus amplification were performed as described for *Hs*ORC^1–5^. The coding sequence of truncated Cdt1 (Cdt1ΔN, amino acids 167-546) was cloned as N-terminal 6xHis-TEV fusion into a LIC-converted, pET-derived *E. coli* expression vector (Macrolab, University of California Berkeley, USA).

*Hs*MCM2-7 was reconstituted in insect cells. MCM2-7 subunits were cloned into two pFastBac-derived BioBricks MultiBac expression vectors^71^, one construct containing Mcm2 (with or without a C-terminal msfGFP tag), Mcm4 (natural variant L650M), and Mcm6, and the other Mcm3, Mcm5, and Mcm7. For affinity purification, Mcm3 and Mcm4, or Mcm5 and Mcm7, were tagged N-terminally with MBP and 6xHis, respectively, each followed by a TEV protease cleavage site. Full-length *Hs*Geminin was cloned into a LIC-converted, pET-derived vector *E. coli* expression (Macrolab, University of California Berkeley, USA) with an N-terminal 6xHis-TEV tag.

### Expression and purification of recombinant *Hs*ORC

Full-length or Orc1-truncated *Hs*ORC^1–5^ and *Hs*ORC^2–5^ were purified at 4°C from Hi5 cells infected with baculoviruses expressing a combination of single or multiple *Hs*ORC subunits. After 48 hours infection, Hi5 cells were harvested and resuspended in 35 mL lysis buffer (50 mM Tris-HCl (pH 7.8), 300 mM KCl, 50 mM imidazole, 10% glycerol, 200 μM PMSF, 1 μg/mL leupeptin, 1 mM β-ME) per liter culture. The cell suspension was sonicated and the lysate clarified by ultracentrifugation at 142,032 × g for 45 min in a Beckman Coulter Optima L-80 XP ultracentrifuge. The soluble fraction was subjected to ammonium sulfate precipitation (final 20% (v/v)) and subsequently cleared once more by ultracentrifugation. The supernatant was loaded onto a 5 mL HisTrap HP Nickel-affinity chromatography column (Cytiva) that was washed with 60 mL lysis buffer prior to *Hs*ORC elution with a 50-250 mM imidazole gradient. Peak fractions were further purified on 7.5-10 mL amylose columns (New England Biolabs) in 50 mM Tris-HCl (pH 7.8), 300 mM KCl, 10% glycerol, 1 mM β-ME and eluted with 20 mM maltose. *Hs*ORC was incubated with 6xHis-tagged TEV protease overnight, which was removed by nickel-affinity chromatography using a 5 mL HisTrap HP column (Cytiva, equilibrated in 50 mM Tris-HCl (pH 7.8), 300 mM KCl, 10% glycerol, 50 mM imidazole, 1 mM β-ME). The flow-through was concentrated and loaded onto HiPrep 16/60 Sephacryl S-400 HR or Superose 6 Increase 10/300 GL columns (Cytiva) equilibrated in 25 mM HEPES-KOH (pH 7.6), 500 mM potassium glutamate, 10% glycerol, 1 mM DTT. *Hs*ORC peak fractions were pooled and concentrated in 30 K Amicon Ultra-15 concentrators (Millipore), aliquoted, and flash-frozen in liquid nitrogen for storage at -80 °C.

### Expression and purification of recombinant *Hs*Orc6

All Orc6 constructs were expressed in BL21 RIL *E. coli* cells. 2 liters of shaker culture in 2xYT media with 30 μg/mL kanamycin and 34 μg/mL chloramphenicol were grown at 37°C to an OD_600 nm_ of 0.5-0.6. For Orc6 mutants, the media was supplemented with 2% glucose. Expression was induced with 0.5 mM Isopropyl β-D-1-thiogalactopyranoside (IPTG) for ∼18 hours at 16°C. Cells were harvested by centrifugation and lysed by sonication in 50 mL lysis buffer containing 50 mM Tris-HCl (pH 8.0), 800 mM KCl, 10% glycerol, 30 mM imidazole, 1 mM β-ME, 200 μM PMSF, and 1 μg/mL leupeptin. The lysate was clarified by centrifugation at 23,426 x *g* for 35 minutes and loaded on a 5 mL HisTrap HP column (Cytiva). After a wash with 200 mL lysis buffer, Orc6 was eluted with 50 mM Tris-HCl (pH 8.0), 150 mM KCl, 250 mM imidazole, 10% glycerol, 1 mM β-ME. The 6xHis tag was cleaved by digestion with 6xHis-tagged TEV protease during overnight dialysis into 50 mM Tris-HCl (pH 8.0), 150 mM KCl, 10% glycerol, 30 mM imidazole, and 1 mM β-ME. TEV protease and uncleaved Orc6 were removed by passing the protein solution over a 5 mL HisTrap HP column (Cytiva) equilibrated in dialysis buffer. Subsequently, cleaved Orc6 was further purified by gel filtration chromatography on a HiLoad 16/60 Superdex 75 pg column (Cytiva) equilibrated in 50 mM Tris-HCl (pH 8.0), 150 mM KCl, 10% glycerol, 1 mM DTT. Peak fractions were pooled, concentrated, aliquoted, and flash-frozen in liquid nitrogen for storage at -80°C.

### Expression and purification of recombinant *Hs*Cdc6

Full-length *Hs*Cdc6 was purified at 4°C from Hi5 cells. After 48 hours infection, Hi5 cells were harvested, resuspended in 35 mL lysis buffer (50 mM Tris-HCl (pH 7.8), 300 mM KCl, 30-50 mM imidazole, 10% glycerol, 200 μM PMSF, 1 μg/mL leupeptin, 1 mM β-ME) per liter culture, and sonicated. The lysate was then clarified by two rounds of ultracentrifugation and ammonium sulfate precipitation. The supernatant was loaded onto a 5 mL HisTrap HP Nickel-affinity chromatography column (Cytiva) that was washed with 60 mL lysis buffer prior to elution with 250 mM imidazole. The protein was further purified on a 5-10 mL amylose column (New England Biolabs) in 50 mM Tris-HCl (pH 7.8), 300 mM KCl, 10% glycerol, 1 mM β-ME and eluted with 20 mM maltose. The *Hs*Cdc6 elution was concentrated in a 30 K Amicon Ultra-15 concentrator (Millipore) and loaded onto a HiLoad 16/600 Superdex 200 pg column (Cytiva) equilibrated in 50 mM Tris-HCl (pH 7.8), 300 mM KCl, 10% glycerol, 1 mM DTT. Peak fractions were incubated with 6xHis-tagged TEV protease overnight and then purified over another 5 mL HisTrap HP column (Cytiva) equilibrated in 50 mM Tris-HCl (pH 7.8), 300 mM KCl, 10% glycerol, 50 mM imidazole, 1 mM β-ME) to remove TEV, uncleaved Cdc6, and 6xHis-MBP. A final gel filtration chromatography step was performed using a HiLoad 16/600 Superdex 200 pg column (Cytiva). Afterwards, *Hs*Cdc6 peak fractions were pooled, concentrated, aliquoted, and flash-frozen in liquid nitrogen for storage at -80°C. Truncated *Hs*Cdc6 was purified similarly to the full-length protein except that the amylose affinity step was omitted.

### Expression and purification of recombinant *Hs*Cdt1

Full-length *Hs*Cdt1 was expressed and purified at 4°C from Hi5 cells infected with baculovirus for 48 hours. Cells were harvested and resuspended in 60 mL lysis buffer (50 mM Tris-HCl (pH 7.8), 1 M NaCl, 10% glycerol, 30 mM imidazole, 5 mM β-ME, 200 μM PMSF, 1 μg/mL leupeptin) per liter culture. After sonication, the lysate was clarified by two rounds of ultracentrifugation and ammonium sulfate precipitation as described for *Hs*ORC. The supernatant was loaded onto a 5 mL HisTrap HP Nickel-affinity chromatography column (Cytiva) that was washed with 200 mL lysis buffer and 50 mL low salt buffer (50 mM Tris-HCl (pH 7.8), 300 mM KCl, 10% glycerol, 30 mM imidazole, 5 mM β-ME), eluted onto a 5 mL HiTrap Q HP ion exchange column (Cytiva) with 250 mM imidazole in 50 mM Tris-HCl (pH 7.8), 300 mM KCl, 10% glycerol, 5 mM β-ME. Fractions containing Cdt1 were further purified on a 5 mL amylose column (New England Biolabs) in 50 mM Tris-HCl (pH 7.8), 300 mM KCl, 10% glycerol, 5 mM β-ME and eluted with 20 mM maltose. The 6xHis-MBP-tag was cleaved off by addition of 6xHis-tagged TEV protease overnight, after which TEV was removed using a HisTrap HP column (Cytiva). The flow-through was concentrated and loaded onto a HiLoad 16/600 Superdex 200 pg column (Cytiva) equilibrated in 50 mM Tris-HCl (pH 7.8), 150 mM KCl, 10% glycerol, 1 mM DTT. *Hs*Cdt1 peak fractions were pooled and concentrated in a 30 K Amicon Ultra-15 concentrator (Millipore), aliquoted, and flash-frozen in liquid nitrogen for storage at - 80°C.

Truncated *Hs*Cdt1 (amino acids 167-546) was expressed in BL21 RIL *E. coli* cells grown in 2xYT media with 30 μg/mL kanamycin and 34 μg/mL chloramphenicol. Expression was induced at an OD_600 nm_ of 0.5-0.6 with 0.5 mM IPTG overnight at 16°C. Cells were harvested by centrifugation and resuspended in ∼20 mL lysis buffer (50 mM Tris-HCl (pH 7.8), 1 M NaCl, 10% glycerol, 30 mM imidazole, 5 mM β-ME, 200 μM PMSF, 1 μg/mL leupeptin) per liter culture. The cell suspension was sonicated and the lysate was clarified by centrifugation at 23,426 × g in a Sorvall Evolution RC Superspeed centrifuge (Thermo Fisher Scientific). Cdt1ΔN was purified by nickel-affinity and ion exchange chromatography as described for the full-length protein. Subsequently, the affinity tag was removed by digestion with 6xHis-tagged TEV protease overnight during dialysis into 50 mM Tris-HCl (pH 7.8), 300 mM KCl, 10% glycerol, 5 mM β-ME, followed by nickel-affinity chromatography to remove TEV and uncleaved Cdt1. The flowthrough was concentrated and loaded onto a HiLoad 16/600 Superdex 200 pg column (Cytiva) equilibrated in 50 mM Tris-HCl (pH 7.8), 150 mM KCl, 10% glycerol, 1 mM DTT. *Hs*Cdt1ΔN peak fractions were pooled and concentrated in a 30 K Amicon Ultra-15 concentrator (Millipore), aliquoted, and flash-frozen in liquid nitrogen for storage at -80°C.

### Expression and purification of recombinant *Hs*MCM2-7

*Hs*MCM2-7 (containing untagged Mcm2 or C-terminally msfGFP-tagged Mcm2) were purified at 4°C from Hi5 cells co-infected with baculoviruses expressing *Hs*Mcm2-Mcm4-Mcm6 and *Hs*Mcm3-Mcm5-Mcm7 subunits. Cells were harvested by centrifugation and resuspended in 35 mL lysis buffer (50 mM HEPES-KOH (pH 7.5), 300 mM potassium acetate, 10% glycerol, 30 mM imidazole, 1 mM β-ME, 1 μg/mL leupeptin, 200 μM PMSF) per liter culture. After sonication, the lysate was clarified by two rounds of ultracentrifugation and ammonium sulfate precipitation as described for *Hs*ORC. *Hs*MCM2-7 was purified using a 5 mL HisTrap HP column (Cytiva) and eluted using 250 mM imidazole in 50 mM HEPES-KOH (pH 7.5), 200 mM potassium acetate, 10% glycerol, 1 mM β-ME. Eluted proteins were loaded onto a 6 mL amylose column (New England Biolabs), washed in 50 mM HEPES-KOH (pH 7.5), 300 mM potassium acetate, 10% glycerol, 1 mM β-ME, and MCM2-7 was eluted with 20 mM maltose in wash buffer. Affinity tags were removed by digestion with 6xHis-tagged TEV protease and passing the protein solution over another 5 mL HisTrap HP column (Cytiva) equilibrated in 50 mM HEPES-KOH (pH 7.5), 200 mM potassium acetate, 30 mM imidazole, 10% glycerol, 1 mM β-ME. The flow-through was concentrated and loaded onto a Superose 6 Increase 10/300 GL column (Cytiva) in 25 mM HEPES-KOH (pH 7.5), 300 mM potassium acetate, 10% glycerol, 1 mM DTT. *Hs*MCM2-7 peak fractions were pooled, concentrated, and flash-frozen in liquid nitrogen for storage at -80°C. For purification of MBP-MCM2-7 (with MBP on Mcm5), the TEV cleavage step was omitted and the complex directly loaded onto a Superose 6 column after amylose affinity chromatography.

### Expression and purification of recombinant *Hs*Geminin

*Hs*Geminin was expressed in *E. coli* BL21 RIL cells overnight at 16°C upon induction with 0.5 mM IPTG at an OD_600 nm_ of 0.5. Cells were harvested, resuspended in 25 mL lysis buffer per liter culture (50 mM Tris-HCl (pH 7.8), 800 mM KCl, 50 mM imidazole, 10% glycerol, 1 mM β-ME, 1 μg/mL leupeptin, 200 μM PMSF), and lysed by sonication. The clarified lysate was used to purify *Hs*Geminin by nickel-affinity chromatography on a 5 mL HisTrap HP column (Cytiva), which was washed with 350 mL lysis buffer. *Hs*Geminin was eluted with 50 mM Tris-HCl (pH 7.8), 600 mM KCl, 250 mM imidazole, 10% glycerol, 1 mM β-ME. After concentration of the eluate, the protein was further purified by size exclusion chromatography in 50 mM Tris-HCl (pH 7.8), 300 mM KCl, 10% glycerol, 1 mM DTT using a HiLoad 16/600 Superdex 200 pg column (Cytiva). Protein in peak fractions was concentrated, flash frozen, and stored at -80°C.

### MCM2-7 loading assay

*In vitro* MCM2-7 loading assays using human proteins were adapted from those established previously for the *Drosophila* system^32^. 2.4 pmol of 178 bp biotinylated duplex DNA (described in reference 32) were coupled to 10 μL streptavidin sepharose high performance beads (Cytiva) in ∼20 μL coupling buffer (5 mM Tris-HCl (pH 8.0), 0.5 mM EDTA, 1 M NaCl, 0.01% NP-40) for 30 minutes. The beads were washed with coupling buffer and 2.4 pmol streptavidin (Sigma Aldrich) were added for 15 minutes to block any free DNA ends. Subsequently, beads were washed thrice in low salt buffer (25 mM HEPES-KOH (pH 7.6), 300 mM potassium glutamate, 10 mM magnesium acetate, 10% glycerol, 0.01% NP-40, 1 mM DTT). Full-length or truncated ORC, Cdc6, Cdt1, and MCM2-7 were mixed at 60 nM, 60 nM, 120 nM, and 120 nM, respectively, as the standard final concentration in low-salt buffer with or without 1 mM ATP or ATPγS. In assays with Geminin, ORC, Cdc6, Cdt1, and Mcm2-7 were mixed with Geminin prior to addition to DNA-bound beads. 40 μL of the protein mix were added to DNA-coupled beads and incubated for 30 minutes at 37°C. To assess MCM2-7 loading, beads were washed once with high-salt buffer (25 mM HEPES-KOH (pH 7.6), 1 M KCl, 10 mM magnesium acetate, 10% glycerol, 0.01% NP-40, 1 mM DTT) with 0 or 1 mM ATP, followed by two washes with low-salt buffer with 0 or 1 mM ATP. To assess MCM2-7 recruitment, beads were washed thrice with low-salt buffer. Bound proteins were eluted by digestion with 500 units micrococcal nuclease (New England Biolabs) for 10 minutes at 37°C in 25 mM HEPES-KOH (pH 7.6), 300 mM KCl, 10 mM magnesium acetate, 10% glycerol, 0.01% NP-40, 5 mM CaCl_2_, 1 mM DTT, 0 or 1 mM ATP (for loading) or 1 mM ATPγS (for recruitment), and analyzed by SDS-PAGE and silver staining. All loading assays were performed at least as three independent experiments. For quantification of Geminin titrations, the band intensity of Mcm2 was measured in ImageJ and normalized to the Mcm2 intensity from recruitment or loading reactions done without Geminin for each experiment. Generally, all assays were performed with truncated Orc1 and Cdt1 unless stated otherwise.

Fluorescence-based MCM2-7 loading assays were performed similarly with the following modifications. 3 pmol of biotinylated DNA were coupled to 10 μL Dynabeads MyOne Streptavidin T1 (Thermo Fisher Scientific) and free ends blocked with 3 pmol streptavidin afterwards. MCM2-7 recruitment and loading reactions were performed at 37°C for 40 minutes in 40 μL each containing 60 nM ORC^1–5^ (with Orc1ΔN), 60 nM Cdc6, 120 nM Cdt1ΔN, and 120 nM GFP-MCM2-7. Beads were washed once each with high-salt wash and low-salt wash buffer for loading reactions, and twice with low-salt wash buffer for recruitment reactions. MNase elutions were transferred to a 384-well plate and GFP fluorescence in eluates measured in a PHERAstar FSX plate reader (BMG Labtech) with excitation at 485 nm and emission at 520 nm using a gain setting of 700. Fluorescence intensities from at least three independent experiments were normalized to the average GFP signal from recruitment or loading reactions done with wildtype proteins. For ORC/Cdc6 titrations, ORC^1–5^ (Orc1ΔN), Orc6, and Cdc6 concentrations of 30, 60, 120, 240, and 480 nM were used while GFP-MCM2-7 and Cdt1ΔN were kept constant at 120 nM. GFP fluorescence in eluates was measured with a gain setting of 500 and 700 for recruitment and loading, respectively. Background fluorescence was accounted for by subtracting the fluorescence intensity measured from a reaction without ATP/ATPγS. Relative MCM2-7 loading efficiencies were calculated by dividing the measured GFP fluorescence intensities in eluates of reactions without Orc6 by those with Orc6 for at least three independent replicate experiments.

To assess dimerization of independently loaded MCM hexamers, MCM was first loaded onto bead-coupled DNA in 40 μL reactions containing 120 nM of ORC^1–5^ (Orc1ΔN), Cdc6, Cdt1ΔN, and MBP-MCM2-7 (MBP-Mcm5) for 30 minutes at 37°C. Due to the lower stability of loaded single hexamers, beads were washed twice with a modified high-salt buffer using 0.5 M KCl instead of 1 M KCl, and once with low-salt buffer (with 1 mM ATP). Beads were then resuspended in 40 μL low-salt buffer (with 1 mM ATP) with or without 600 nM Orc6 or Geminin and incubated for 10 minutes at 30°C. Subsequently, TEV protease was added to a final concentration of 2.5 μM, and reactions were incubated for 30 minutes at 30°C and another 10 minutes at 37°C. Beads were washed with high-salt (0.5 M KCl) and low-salt buffer with 1 mM ATP. MCM2-7 was eluted by MNase digestion as described above and 4 μL of the elution were used to prepare negative-stain EM grids (see below). Functionality of cleaved MCM2-7 purified with MBP-Mcm5 was confirmed in standard loading assays by pre-treating MCM2-7 with 5 μM TEV for 30 minutes at 30°C.

### Negative-stain EM

Samples for negative staining were prepared as described below, applied to glow-discharged, continuous carbon EM grids (Ted Pella), and stained with 2% uranyl acetate. EM grids were imaged in a Talos L120C transmission electron microscope operated at 120 kV. Micrographs were either collected manually or automatically using serial EM^72^ at magnifications of 36 kx, 45 kx, or 73 kx.

#### MCM2-7 loading reaction eluates

To analyze the outcome of bead-based MCM2-7 loading assays by negative-stain EM, 4 μL of proteins eluted from streptavidin beads were applied to glow-discharged continuous carbon grids for 1 minute (or 5 minutes for MCM dimerization assays from loaded single hexamers) and stained with uranyl acetate. MCM2-7 double hexamer particles per micrograph were counted manually. A total of three independent replicate experiments were done for each experimental condition, with a minimum of 10 micrographs analyzed per replicate. The data were normalized to the average number of MCM2-7 double hexamers per micrograph observed in reactions containing all loading factors (full-length or wildtype proteins). Statistical significance was calculated using two-way ANOVA with Tukey post-hoc analysis in GraphPad Prism.

#### MCM2-7 loading intermediates

ORC^1–5^ (Orc1ΔN), Cdc6, Cdt1ΔN, and MCM2-7 were mixed at 120 nM each with 180 nM biotinylated DNA and 360 nM streptavidin, either with or without 120 nM Orc6, in low-salt buffer (25 mM HEPES-KOH (pH 7.6), 300 mM potassium glutamate, 10 mM magnesium acetate, 10% glycerol, 0.01% NP-40, 1 mM DTT, 1 mM ATP or ATPγS). To simplify interpretation of intermediates, MCM2-7 was incubated at 37°C for 20 minutes prior to assembling loading reactions to dissociate MCM2-7 dimers. Loading reactions were incubated at 37°C for 30 minutes (or shorter for time course experiments). Subsequently, 4 μL of loading reactions were used for preparing EM grids. Typically, two datasets from independently prepared loading reactions were collected for each condition at 45 kx magnification, with approximately 150 micrographs for each dataset. Particles were picked automatically (reference-free) using GAUTOMATCH^73^, extracted from phase-flipped micrographs (done using GCTF^74^), and subjected to 2D classification in RELION 4.0.1^75^.

For structural analysis of MO intermediates, MO-like particles from all datasets were combined and used for *ab-initio* reconstruction in CryoSPARC v4.2.1^76^. Two classes were identified that contained clear density for an MCM hexamer but differed in the orientation of ORC. Both volumes were refined using one round of heterogeneous and homogeneous refinement in CryoSPARC. Docking of MCM-SH (this study), a single hexamer from the loaded MCM-DH (PDB 7w1y^29^), and ORC (PDB 7jpo^77^) into the EM maps was done using UCSF Chimera^78,79^. The orientation of MCM2-7 was determined using the winged helix domains Mcm2, Mcm5, and Mcm6 as reference point because they interact and form a prominent map region at the C-terminal side of the MCM2-7 ring^29^. 2D projections for comparison with 2D class averages were generated by converting model coordinates into maps and low-pass filtering volumes to 25 Å.

#### Purified HsMCM2-7 (pre-loading state)

Purified, recombinant MCM2-7 was diluted to 80 nM in 25 mM HEPES-KOH (pH 7.5), 10 mM magnesium acetate, 0.3 M potassium glutamate, 10% glycerol, 1 mM DTT, either in the absence or presence of 1 mM ATP and/or 240 nM full-length Cdt1 (final concentration). 4 μL of each sample were applied to EM grids prior to staining. Equilibrium studies were performed at 2 μM MCM2-7 and the complex was diluted to 80 nM prior to EM grid preparation.

### Cryo-EM of MCM2-7 loading reactions

#### Cryo-EM sample preparation and data collection

Truncated ORC^1–5^ (Orc1ΔN), Orc6, 6xHis-MBP-Cdc6, and a 480 bp dsDNA fragment (obtained by PCR of the *S. cerevisiae* ARS1 locus with oligos 5’-GGACTGACGCCAGAAAATGTTG-3’ and 5’-CGAGGATACGGAGAGAGGTATG-3’) were mixed at 100 nM concentration with 200 nM truncated Cdt1 and MCM2-7 in 25 mM HEPES-KOH (pH 7.6), 10 mM magnesium acetate, 0.3 M potassium glutamate, 1 mM DTT, and 1 mM ATP. MCM2-7 was preincubated at 37°C for 20 minutes to disassemble MCM2-7 dimers prior to mixing with the other proteins. After a 5-minute incubation at 37°C, 3.5 μL of this protein-DNA mix were applied to 300-mesh copper R2/1 Quantifoil Holey Carbon Grids (Quantifoil Micro Tools GmbH) that had been freshly glow discharged for 30 seconds at 25 mA in a GloQube Glow Discharge System (Quorum Technologies) and treated with graphene oxide (Sigma Aldrich) as described previously^80^. Grids were blotted for 4 seconds and vitrified in liquid ethane using a Vitrobot Mark IV plunge freezer (Thermo Fisher Scientific). Cryo-EM grids were pre-screened in a 200 kV Glacios transmission electron microscope. Cryo-EM data were collected in two separate sessions on a Titan Krios G2 300 kV transmission electron microscope (Thermo Fisher Scientific) with a post-GIF K3 summit direct electron detector (Gatan) using SerialEM^72^. The target defocus was set to -1.0 to -2.0 µm. 5 exposures per 2 μm hole were recorded as movies with 40 frames in super-resolution mode at a physical pixel size of 0.832 Å for a total of ∼1.4 seconds with a dose rate of 34.7-36.1 e^-^/Å^2^ per second, yielding a total electron dose of 50-51 e^-^/Å^2^.

#### Cryo-EM data processing and model building of loaded MCM single and double hexamers

All processing steps were performed using CryoSPARC v4^76,81^. Raw movies were pre-processed using CryoSPARC Live, which included motion correction, Fourier cropping to 0.5, and estimation of CTF parameters. Particles were picked in CryoSPARC v4 using the blob pick tuner function, extracted with a box size of 600 x 600 pixels, and Fourier-cropped to a box size of 300 x 300 pixels, resulting in a pixel size of 1.664 Å. Iterative 2D classification in CryoSPARC v4 was used to remove ‘junk’ particles from each of the two datasets. Subsequently, particles from both datasets were combined and subjected to additional rounds of 2D classification, followed by *ab initio* reconstruction and heterogeneous refinement, which yielded a well-resolved 3D volume for the loaded MCM2-7 double hexamer (closed MCM-DH). Non-MCM-DH particles were further cleaned by 2D classification, which identified a small subset of particles that corresponded to MCM-DH (∼8 K) that were combined with the other MCM-DH particles during subsequent 3D refinement. Non-MCM-DH particles that resembled single MCM2-7 hexamers were used as input for *ab initio* reconstruction and heterogeneous refinement, which yielded a 3D volume of a closed-ring, single MCM2-7 hexamer with duplex DNA inside the central MCM pore and corresponded to DNA-loaded MCM2-7 single hexamers (closed MCM-SH). Following 3D refinement, closed MCM-DH and MCM-SH particles were re-extracted at the original pixel size (0.832 Å) with a box size of 500 x 500 pixels. Final 3D maps were calculated using non-uniform refinement in CryoSPARC v4. CTF refinement improved the resolution of the closed MCM-DH cryo-EM reconstruction but not of that of the closed MCM-SH (and was therefore omitted in this case). Cryo-EM maps were sharpened with DeepEMhancer^82^. Resolution values were calculated using gold-standard Fourier Shell Correlation (FSC) method in CryoSPARC v4 and using 3DFSC^83,84^.

For model building, one or both hexamers from PDB 7w1y^29^ were docked into the cryo-EM map of the loaded MCM-DH and MCM-SH, respectively, using UCSF Chimera^78,79^. The model was checked in COOT^85^ against the cryo-EM map density and minor model rebuilding performed to fix register shift, to account for minor conformational changes between the MCM-SH and MCM-DH structures, and to update the nucleotide status of the ATPase sites of MCM2-7. The models were real-space refined in PHENIX^86,87^ and validated with MolProbity^88^ (**Extended Data Table 1**).

### Structure determination of purified *Hs*MCM2-7 (pre-loading state)

#### Cryo-EM sample preparation and data collection

*Hs*MCM2-7 and truncated *Hs*Cdt1 were mixed at 1.8 μM and 3.6 μM, respectively, in 25 mM HEPES-KOH (pH 7.5), 300 mM potassium acetate, 10 mM magnesium acetate, 10% glycerol, 1 mM ATP, and 1 mM DTT. After a 10-minute incubation on ice, the sample was crosslinked for 10 minutes by adding an equal volume (125 μL) of crosslinking buffer (25 mM HEPES-KOH (pH 7.5), 300 mM potassium acetate, 10 mM magnesium acetate, 10% glycerol, 1 mM DTT, and 0.05% glutaraldehyde). Crosslinking was stopped by the addition of 25 μL quenching solution (900 mM Tris-HCl (pH 7.5), 9 mM lysine, 9 mM aspartate). The crosslinked complex was purified by gel filtration chromatography on a Superose 6 Increase 10/300 GL column (Cytiva) equilibrated in 25 mM HEPES-KOH (pH 7.5), 300 mM potassium acetate, 10 mM magnesium acetate, 0.2 mM ATP, and 1 mM DTT. Peak fractions were concentrated to ∼3.2 mg/mL and 3.5 μL of concentrated sample immediately applied to a Quantifoil R2/1 300 copper mesh grid (Quantifoil Micro Tools GmbH), which had been glow discharged in a GloQube Plus glow discharger for 45 seconds at 25 mA and coated with graphene oxide (Sigma Aldrich) as described in reference 80. The sample was incubated on the grid for 10 seconds, blotted for 3 seconds, and vitrified in liquid ethane using a Vitrobot Mark IV (Thermo Fischer Scientific).

Cryo-EM grids were pre-screened in a 200 kV Glacios transmission electron microscope and then transferred to a 300 kV Titan Krios G2 (Thermo Fischer Scientific) equipped with a post-GIF K3 direct detector (Gatan) for data collection. 40-frame movies (5 movies per hole) were recorded automatically using SerialEM^72^ in super-resolution mode at a nominal magnification of 105,000x and a physical pixel size of 0.832 Å, targeting a defocus range of 1-2 μm and a total electron dose of 51 electrons per Å^2^.

#### Cryo-EM data processing and model building

Super-resolution cryo-EM movies were preprocessed in RELION 4.0, which included motion correction with RELION’s implementation of MotionCor2 and 2-fold binning, as well as estimation of contrast transfer function (CTF) parameters with CTFFIND-4.1^75,89,90^. Particles were picked using RELION’s Laplacian-of-Gaussian filter autopicking implementation, extracted with a box size of 600 x 600 pixels, and downscaled by a factor of 2. Particles were then imported into CryoSPARC v4, which was used for all subsequent processing steps^76^. Particles were first subjected to iterative 2D classification to identify 330,708 good-quality particles that were used for *ab initio* reconstruction and 3D refinement to obtain an initial map at 3.88 Å resolution. To resolve heterogeneity, the particles were further subjected to 3D classification into 8 classes. Particles from the three best 3D classes were kept and combined for another round of 3D classification. One of the two classes displayed good density on both MCM2-7 hexamers, and its particles were re-extracted at the original pixel size prior to non-uniform refinement to yield a final map of the open MCM2-7 dimer that was postprocessed with DeepEMhancer^82^. To improve the local resolution and map quality of a single MCM2-7 hexamer region, we employed C2 symmetry expansion and used this new particle stack for local refinement with a soft-edged mask encompassing one of the MCM2-7 hexamers in the dimer. The resultant 3D volume was post-processed with DeepEMhancer^82^ and used for model building.

For model building, the N- and C-terminal domains of each MCM2-7 subunit from PDB 7w1y^29^ were rigid-body docked into the cryo-EM map using UCSF Chimera^78,79^, manually rebuilt in COOT^85^, and real-space refined in PHENIX^86,87^. To resolve the dimerization interface in the open MCM2-7 dimer, the refined model of the single, open MCM2-7 hexamer was docked into the cryo-EM map of the open MCM2-7 dimer and re-refined in PHENIX. Both refined models were validated using MolProbity^88^ (**Extended Data Table 1**).

To ensure MCM2-7 dimerization was not caused by crosslinking, we also collected a cryo-EM dataset of uncrosslinked MCM2-7 in the absence of Cdt1 and nucleotide. Cryo-EM grids were prepared by applying 3.5 μL of recombinant *Hs*MCM2-7 at 1.5 μM concentration onto a glow-discharged (25 seconds at 25 mA in a GloQube Plus glow discharger) 300-mesh R1.2/1.3 UltrAuFoil grid (Quantifoil Micro Tools GmbH) in 25 mM HEPES-KOH (pH 7.5), 300 mM potassium acetate, 1% glycerol, and 1 mM DTT. Excess sample was removed by blotting for 4 seconds and vitrified as described for the crosslinked sample. Cryo-EM data were collected as done for crosslinked MCM2-7 except that EPU was used for automatic data collection with one exposure per hole and a physical pixel size of 0.825 Å. Data processing was done as described for the crosslinked sample. 2D classification clearly indicated the presence of open MCM2-7 dimers, although their relative orientations were more variable and prevented the reconstruction of high-resolution cryo-EM maps.

## Data availability

The PDB coordinates and cryo-EM maps have been deposited into the Protein Data Bank and Electron Microscopy Data Bank under the following accession numbers: PDB 8W0E and EMD-43707 for the loaded *Hs*MCM2-7 single hexamer, PDB 8W0F and EMD-43708 for the loaded *Hs*MCM2-7 double hexamer, PDB 8W0G and EMD-43709 for *Hs*MCM2-7 dimers, and PDB 8W0I and EMD-43710 for the locally refined map containing one copy of the *Hs*MCM2-7 hexamer from the dimer.

## Acknowledgements

Cryo-EM data were collected at the Yale Cryo-EM resource and at the Laboratory for BioMolecular Structure (LBMS). We would like to thank Shenping Wu and Jianfeng Lin at the Yale Cryo-EM resource, as well as Liguo Wang, Guobin Hu, and Jake Kaminsky at LBMS, for assistance with microscope operation and data collection. We are grateful to Haleigh Marzano for help with initial model building of the MCM dimer. We thank Mark Hochstrasser, Mark Solomon, and members of the Bleichert laboratory for critical reading of the manuscript. This work is supported by the National Institutes of General Medicine (R01-GM141313 to F.B.). O.H. is supported by the NIH predoctoral program in Biophysics (T32-GM008283) and the National Cancer Institute (1F31-CA278331-01). LBMS is supported by the DOE Office of Biological and Environmental Research (KP1607011).

## Author contributions

F.B. conceptualized and supervised the project. R.Y., O.H., M.W., and F.B. cloned expression constructs and purified recombinant proteins. R.Y. and O.H. prepared samples and collected negative-stain EM and cryo-EM data. R.Y., O.H., and F.B. performed biochemical experiments, processed EM data, built and refined atomic models, and wrote the manuscript.

