## Supplemental Data for "Multiple pathways for licensing human replication origins"

### Extended Data Figures

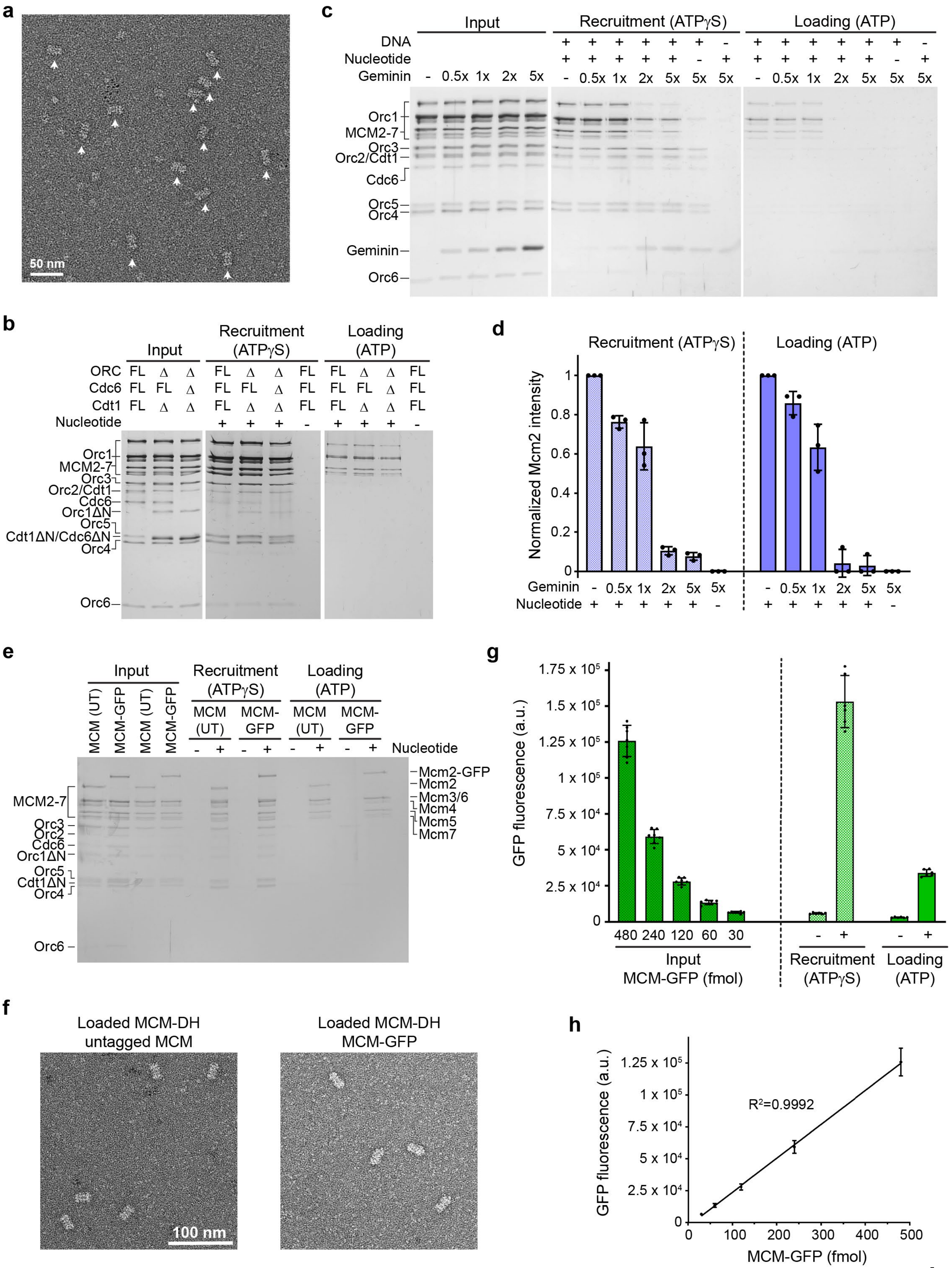
a
50 nm
c
Input
Recruitment (ATPγS)
Loading (ATP)
DNA
Nucleotide
Geminin
- 0.5x 1x 2x 5x - 0.5x 1x 2x 5x 5x 5x - 0.5x 1x 2x 5x 5x 5x
Orc1
MCM2-7
Orc3
Orc2/Cdt1
Cdc6
Orc5
Orc4
Geminin
Orc6
b
Input
Recruitment (ATPγS)
Loading (ATP)
ORC FL Δ Δ FL Δ Δ FL FL Δ Δ FL
Cdc6 FL FL Δ FL FL Δ FL FL FL Δ FL
Cdt1 FL Δ Δ FL Δ Δ FL FL Δ Δ FL
Nucleotide + + + - + + + -
Orc1
MCM2-7
Orc3
Orc2/Cdt1
Cdc6
Orc1ΔN
Orc5
Cdt1ΔN/Cdc6ΔN
Orc4
Orc6
d
Recruitment (ATPγS)
Loading (ATP)
Normalized Mcm2 intensity
1
0.8
0.6
0.4
0.2
0
Geminin - 0.5x 1x 2x 5x 5x - 0.5x 1x 2x 5x 5x
Nucleotide + + + + + - + + + + + -
e
Input
Recruitment (ATPγS)
Loading (ATP)
MCM (UT)
MCM-GFP
MCM (UT)
MCM-GFP
MCM (UT)
MCM-GFP
MCM (UT)
MCM-GFP
- + - + - + - +
Nucleotide
MCM2-7
Orc3
Orc2
Cdc6
Orc1ΔN
Orc5
Cdt1ΔN
Orc4
Orc6
Mcm2-GFP
Mcm2
Mcm3/6
Mcm4
Mcm5
Mcm7
g
GFP fluorescence (a.u.)
1.75 x 10^5
1.5 x 10^5
1.25 x 10^5
1 x 10^5
7.5 x 10^4
5 x 10^4
2.5 x 10^4
0
480 240 120 60 30
Input
MCM-GFP (fmol)
- +
Recruitment (ATPγS)
- +
Loading (ATP)
f
Loaded MCM-DH
untagged MCM
Loaded MCM-DH
MCM-GFP
100 nm
h
GFP fluorescence (a.u.)
1.25 x 10^5
1 x 10^5
7.5 x 10^4
5 x 10^4
2.5 x 10^4
0
R²=0.9992
0 100 200 300 400 500
MCM-GFP (fmol)

**Extended Data Figure 1.** *In vitro* reconstitution of human origin licensing and validation of bead-based MCM loading assays. **a**) Electron micrograph of negatively stained particles from high-salt wash eluate of loading reactions with ATP shows MCM double hexamers (white arrows). **b**) N-terminal, intrinsically disordered regions (IDRs) in Orc1, Cdc6, and Cdt1 are not essential for human MCM loading. Silver-stained SDS-PAGE gels for bead-based MCM recruitment and loading assays are shown. N-terminally truncated Orc1 and Cdt1 were used in all subsequent experiments unless noted otherwise. FL – full-length; Δ – truncated. **c** and **d**) Geminin inhibits MCM recruitment and loading in the *in vitro* reconstituted system. **c**) Silver-stained SDS-PAGE gels of inputs, recruited, and loaded proteins in the absence or presence of increasing concentrations of Geminin (labeled as molar fold excess compared to Cdt1). Note that Geminin associates nonspecifically with beads in low-salt conditions. Full-length ORC and Cdt1 were used in this experiment. **d**) Quantification of results in **c**. Means and standard deviations of Mcm2 band intensities from three independent experiments. Values were normalized to the reaction without geminin on each gel. **e** to **h**) Validation of fluorescence-based MCM loading assay. **e**) MCM2-7 with GFP fused to the C-terminus of Mcm2 supports MCM recruitment and loading to similar levels as untagged (UT) MCM. Silver-stained SDS-PAGE gel is shown. **f**) Loaded MCM-GFP are double hexamers like untagged MCM2-7 as seen in negative-stain EM images of particles eluted from beads. **g**) Raw GFP fluorescence intensities in inputs and elutions from recruitment and loading reactions with MCM-GFP. **h**) The fluorescence intensity signal linearly increases with MCM-GFP concentration. The means and standard deviations from six independent experiments are plotted in **g** and **h**. a.u. – arbitrary units.

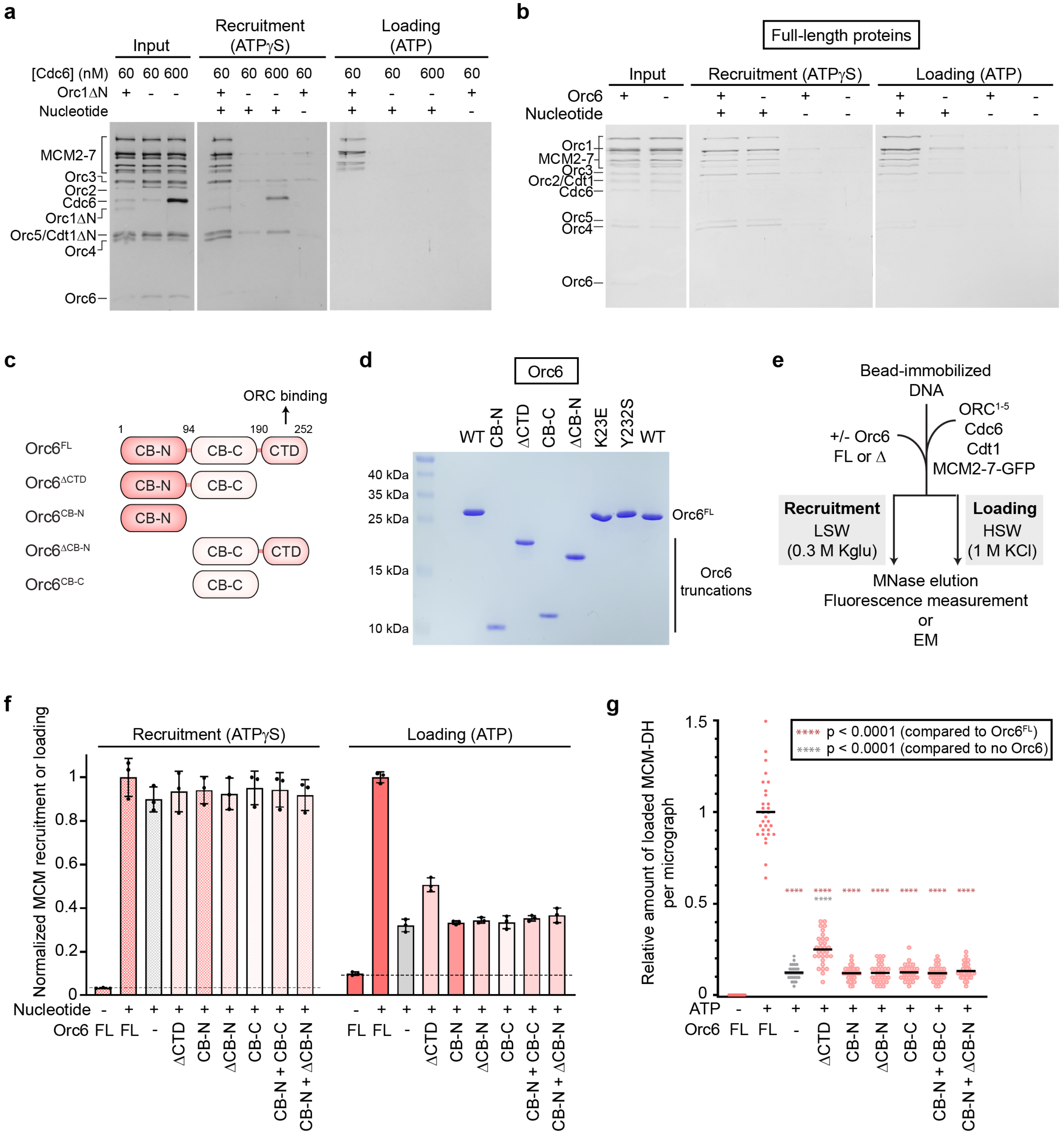


**Extended Data Figure 2.** Full-length Orc6 facilitates while Orc1 is essential for human MCM loading. **a**) Orc1 is required for MCM recruitment (low-salt wash) and loading (high-salt wash) *in vitro* and its absence cannot be complemented by excess Cdc6. Silver-stained SDS-PAGE gels of inputs and elutions from bead-based recruitment and loading reactions are shown. **b**) The IDRs of Orc1 and Cdt1 do not rescue the lower MCM loading efficiency in the absence of Orc6. Silver-stained SDS-PAGE gels of elutions from bead-based recruitment and loading reactions with full-length human loading factors are included. **c** to **g**) All Orc6 domains are required *in cis* to enhance human MCM loading. **c**) Domain architecture of human (*Hs*) Orc6. Numbers denote amino acid positions of domain boundaries. Truncated *Hs*Orc6 constructs used are schematized below. **d**) Coomassie-stained SDS-PAGE gel of purified full-length (FL) wildtype (WT), truncated, and mutant Orc6 proteins used in this study. **e**) Experimental setup for assessing the contributions of Orc6 domains to MCM loading. **f**) MCM loading but not recruitment is impaired when Orc6 is absent or truncated. The means and standard deviations of MCM-GFP fluorescence in elutions of bead-based recruitment and loading assays (normalized to the average signal obtained with full-length (FL) Orc6) from three independent experiments are plotted. Dashed lines mark background signal in recruitment and loading reactions without nucleotide. **g**) Quantification of MCM double hexamers observed by EM in elutions of MCM loading reactions (after high-salt wash) without Orc6 (-), or with full-length (FL) or truncated Orc6. The numbers of double hexamers per micrograph were counted manually from three independent experiments, with 10 micrographs recorded for each sample per experimental repeat (total n=30 micrographs per sample), and normalized to the +Orc6 FL sample. Black lines in the scatter plots represent means. Statistical significance was determined using two-way ANOVA analysis and the Tukey's multiple comparisons test. Note that the differences in MCM loading efficiencies without Orc6 and with Orc6 truncations between **f** and **g** are likely due to the presence of loaded MCM single hexamers that contribute to the signal measured in the fluorescence-based assay in addition to MCM double hexamers, thus slightly overestimating MCM loading efficiency.

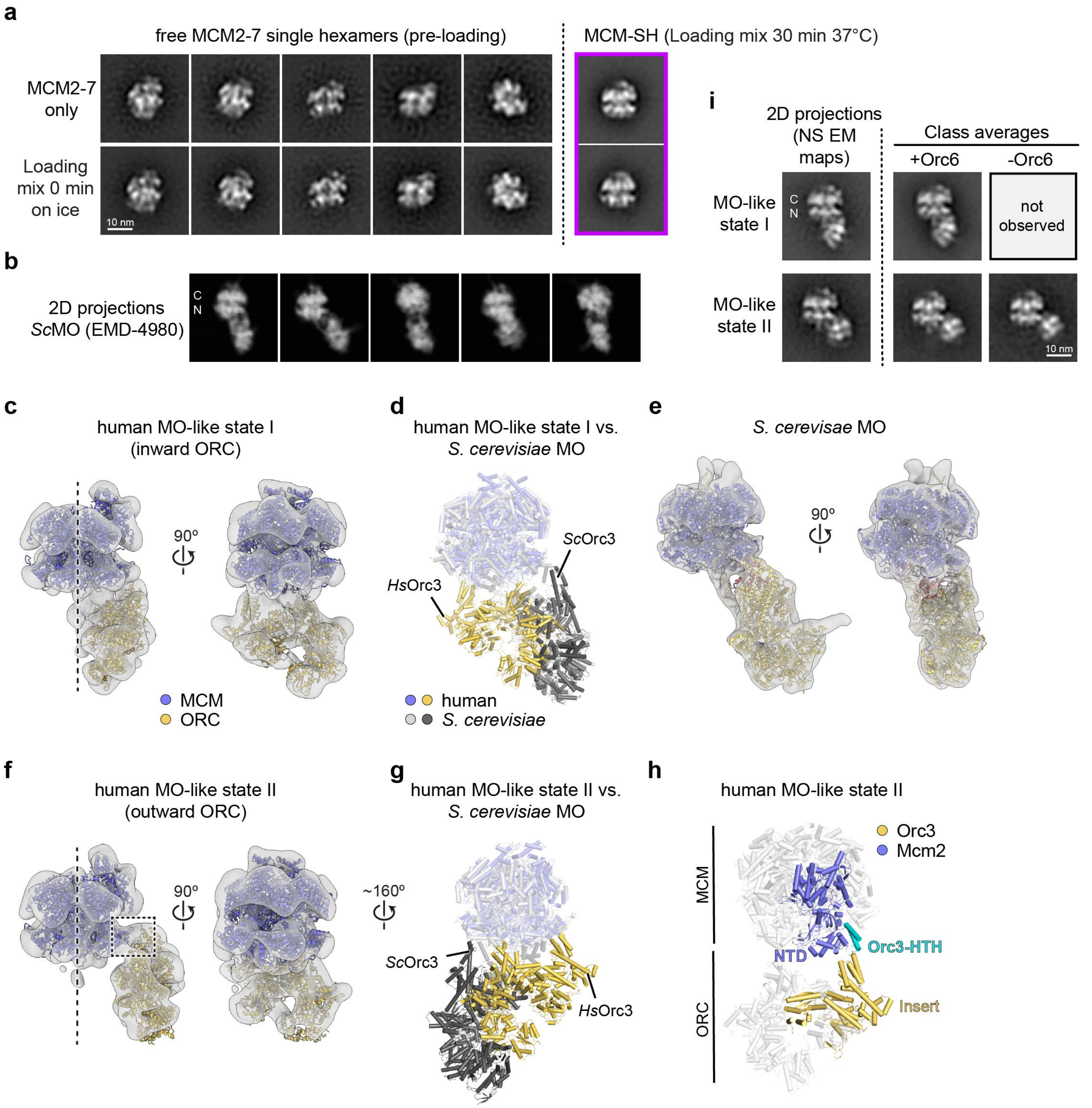


**Extended Data Figure 3.** The human MO-like complex adopts different conformational states that are architecturally distinct from the *S. cerevisiae* MO. **a**) MCM-SH class averages (negative stain) are not observed with purified human MCM2-7 alone or in loading reactions kept on ice, indicating they resemble a conformation distinct from that adopted by free MCM2-7 hexamers. **b**) Projections of the *S. cerevisiae* (*Sc*) MO cryo-EM map (EMD-4980[20], low-pass filtered to 20 Å). Comparison with class averages in Fig. 3c suggests distinct interactions between MCM and ORC in the human and budding yeast complexes. **c** to **g**) Comparison of human MO-like and *S. cerevisiae* MO complex architectures. **c** and **f**) Negative-stain EM reconstructions of human MO-like complexes in states I (in **c**) and II (in **f**). The MCM hexamer structures of human

MCM-SH (this study) or MCM-DH (PDB 7w1y[29]) and human ORC (PDB 7jpo[77]) were docked into negative-stain EM reconstructions. The MCM register was established unambiguously using the winged helix domains of Mcm2, Mcm5, and Mcm6 as a fiducial. **e**) Low-pass filtered (to 15 Å) cryo-EM map of the *S. cerevisiae* MO (EMD-4980 with PDB 6rqc[20]). EM maps in **c**, **e**, and **f** are shown with MCM in the same orientation for easy comparison. Dashed lines in **c** and **f** mark the central axis of MCM. **d** and **g**) Superpositions of MCM in *S. cerevisiae* MO with that in the docked models for human MO-like complexes demonstrate that human ORC engages MCM in a completely distinct manner from *S. cerevisiae* ORC. The Orc6-CTD-binding region in the Orc3 inserts of human and *Sc*ORC are marked for reference. **h**) The Orc3 insert and the N-terminal domain of Mcm2 are positioned in close proximity after docking ORC and MCM into the map of the human state II MO-like complex. This assignment agrees with AlphaFold Multimer prediction of an interaction between a helix-turn-helix (HTH) motif in the human Orc3 insert and the Mcm2-NTD[57,58]. The dashed box in **f** delineates this interaction site in the EM map. **i**) Class averages (negative stain) of human MO-like intermediates in loading reactions with and without Orc6 compared to 2D projections of negative-stain (NS) EM maps of human MO-like states I and II. The class average characteristic of state I is not observed if Orc6 is omitted, whereas those corresponding to state II are seen in the presence and absence of Orc6.

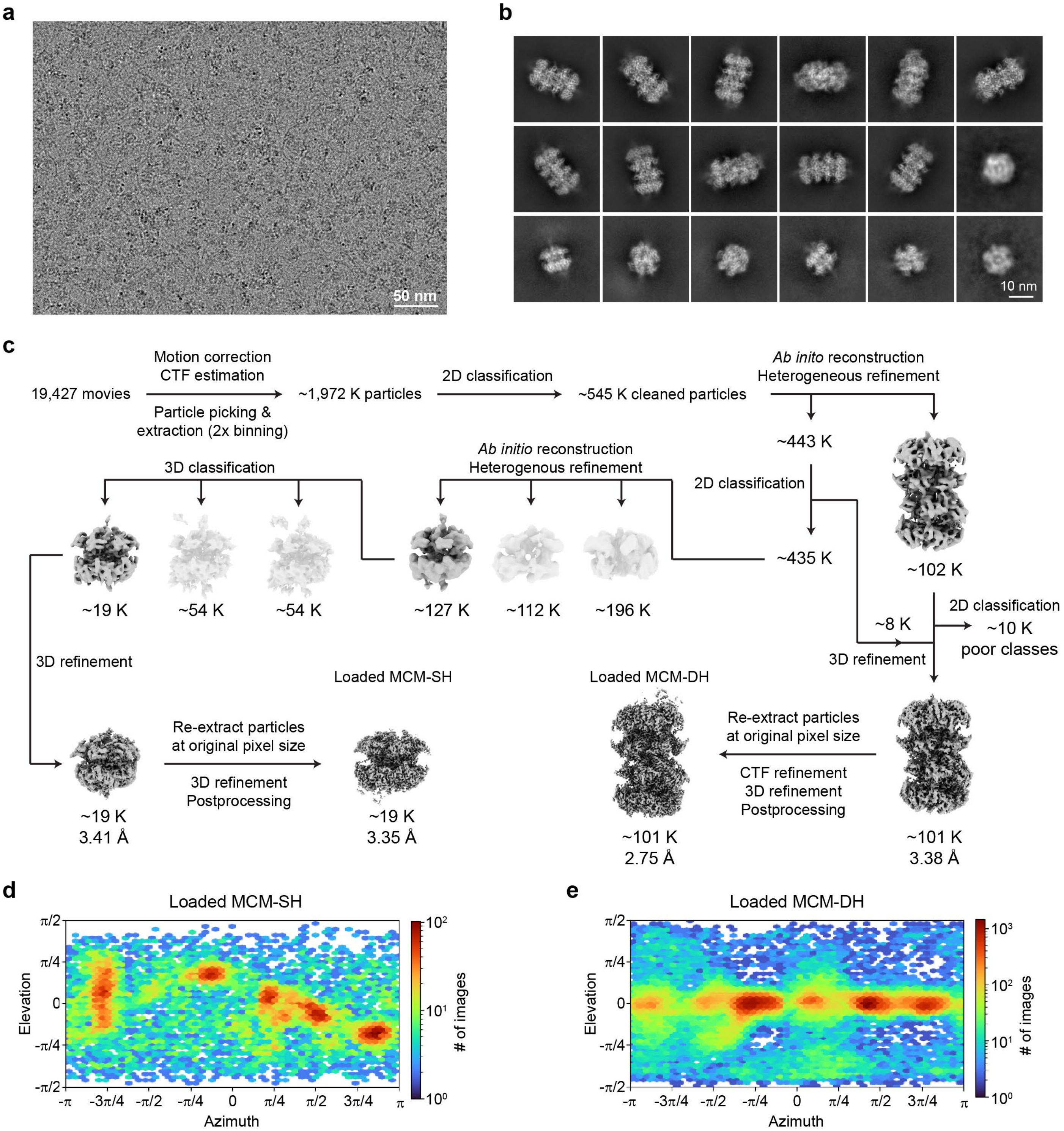


**Extended Data Figure 4.** Cryo-EM data processing and validation of loaded human MCM2-7. **a**) Cryo-EM image and **b**) 2D cryo-class averages from human MCM loading reactions. **c**) Cryo-EM data processing workflow for MCM single and double hexamer reconstructions. **d** and **e**) Angular distribution plots for reconstructed cryo-EM volumes of loaded single (in **d**) and double (in **e**) MCM hexamers.

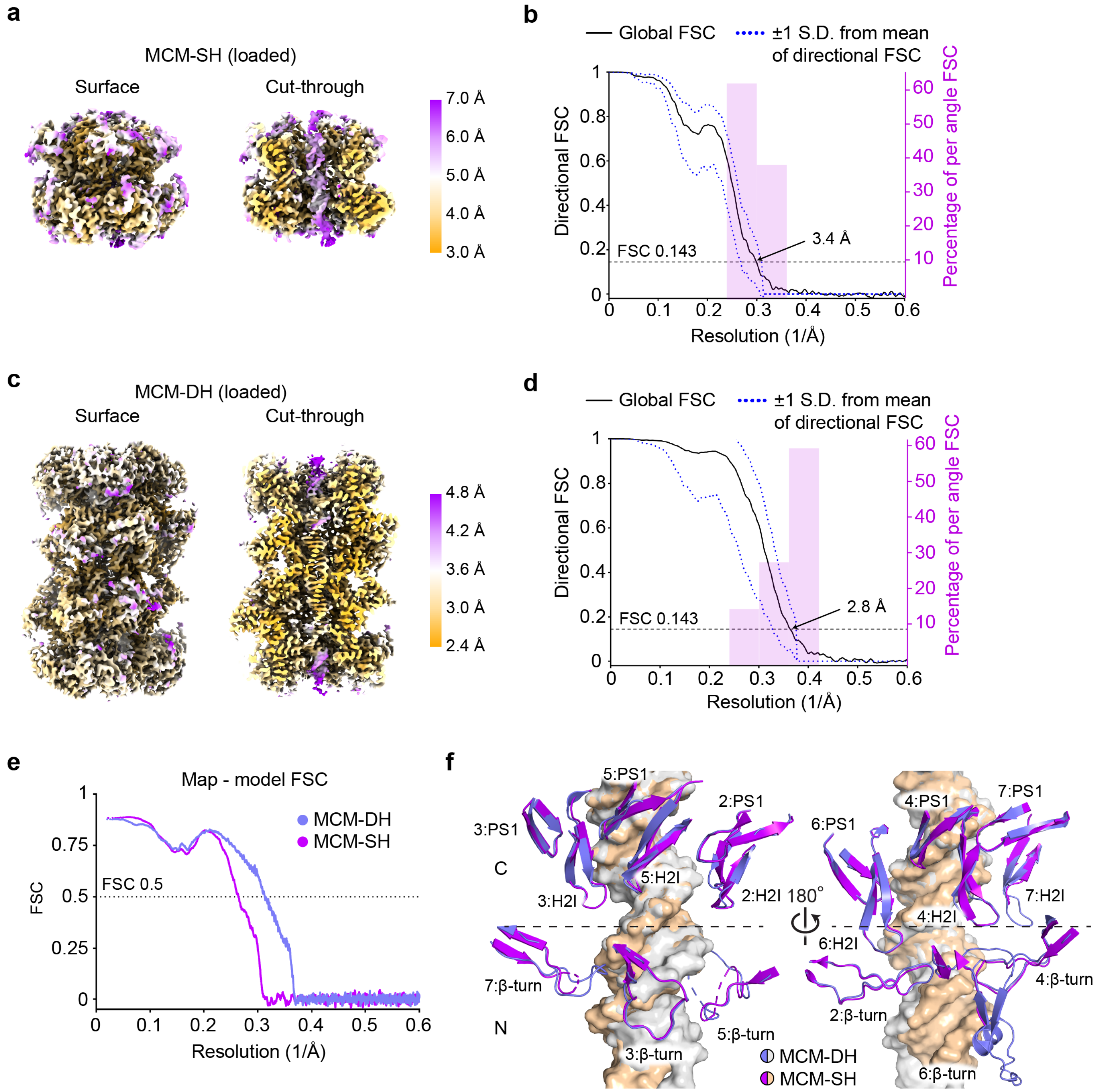


**Extended Data Figure 5.** Resolution estimation of loaded human MCM-SH and MCM-DH cryo-EM reconstructions. **a** and **c**) Surface and cut-through views of unsharpened cryo-EM maps of loaded MCM single (in **a**) and double (in **c**) hexamers. **b** and **d**) 3D Fourier shell correlation (FSC) plots for loaded MCM single (in **b**) and double (in **d**) hexamer cryo-3D reconstructions. **e**) FSC curves of cryo-EM maps and refined models. **f**) Interior hairpin loop interactions with DNA are similar in MCM-DH and MCM-SH. Helix-2-insert (H2I), pre-sensor 1 β hairpin (PS1), and β-turn loops in each subunit are shown as cartoon and DNA is shown as surface. The β-turn loops in Mcm5 of both MCM-DH and MCM-SH, the H2I and β-turn loops in Mcm7 of MCM-SH, and the β-turn loop in Mcm6 of MCM-SH are partially or mostly disordered.

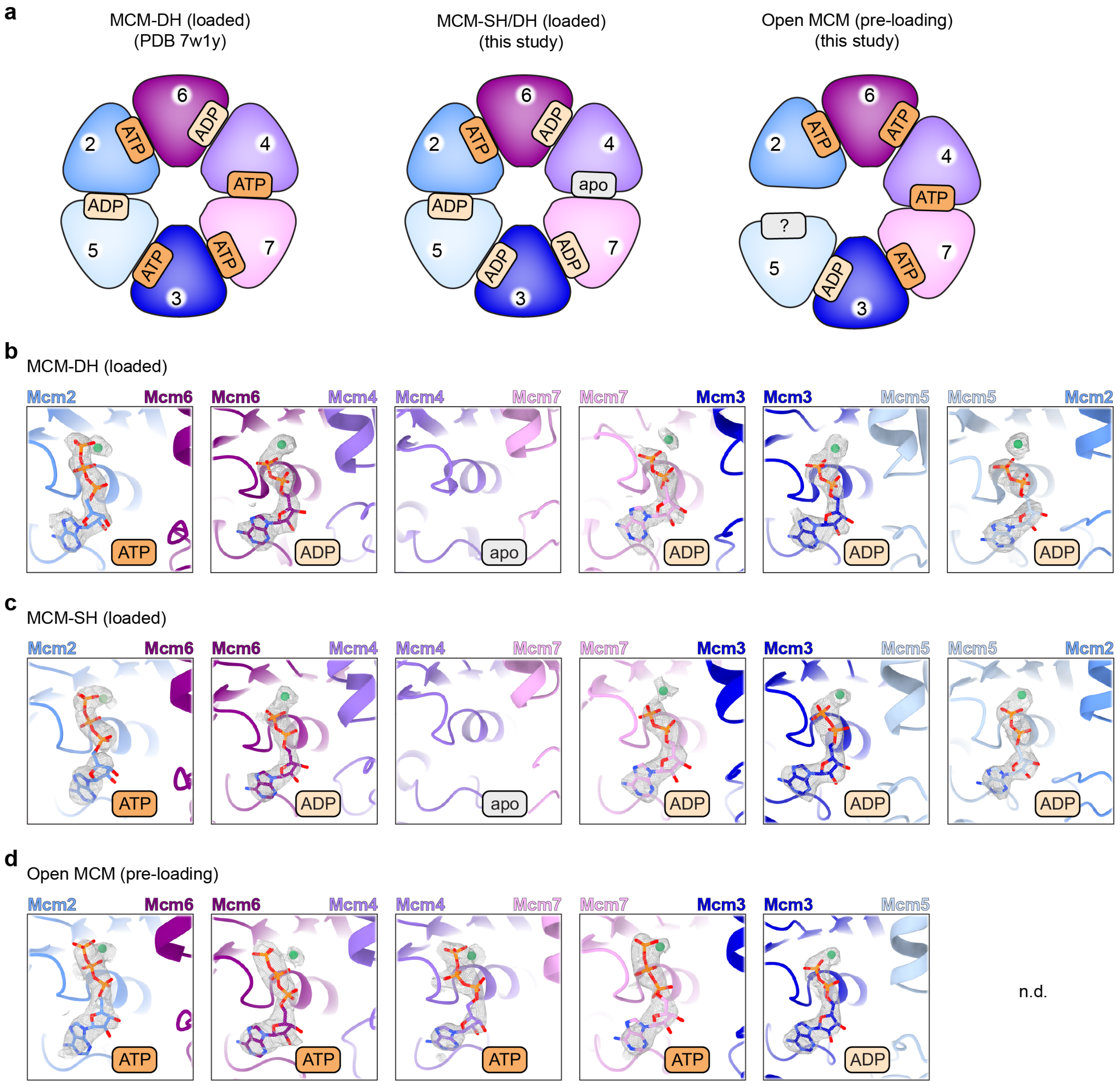


**Extended Data Figure 6.** Nucleotide-binding states within human MCM2-7 before and after loading onto DNA. **a**) Comparison of nucleotide occupancies at the AAA+ interfaces in the loaded MCM single and double hexamers and the open MCM2-7 dimer (pre-loading state) observed in this study with those in the previously published human MCM double hexamer structure (PDB 7w1y[29]). ATPase sites are viewed from the N-terminal MCM tier. **b** to **d**) Zoomed views of the ATP binding sites in the loaded MCM double hexamer (in **b**), the loaded MCM2-7 single hexamer (in **c**), and open-gate MCM2-7 dimer in the pre-loading state (in **d**). ATP and magnesium are shown in stick representation and as spheres, respectively, with corresponding cryo-EM map density as grey mesh. Note that the nucleotide occupancy at the Mcm2-5 site in open MCM2-7 (in **d**) could not be determined (n.d.) due to poor map quality in this region.

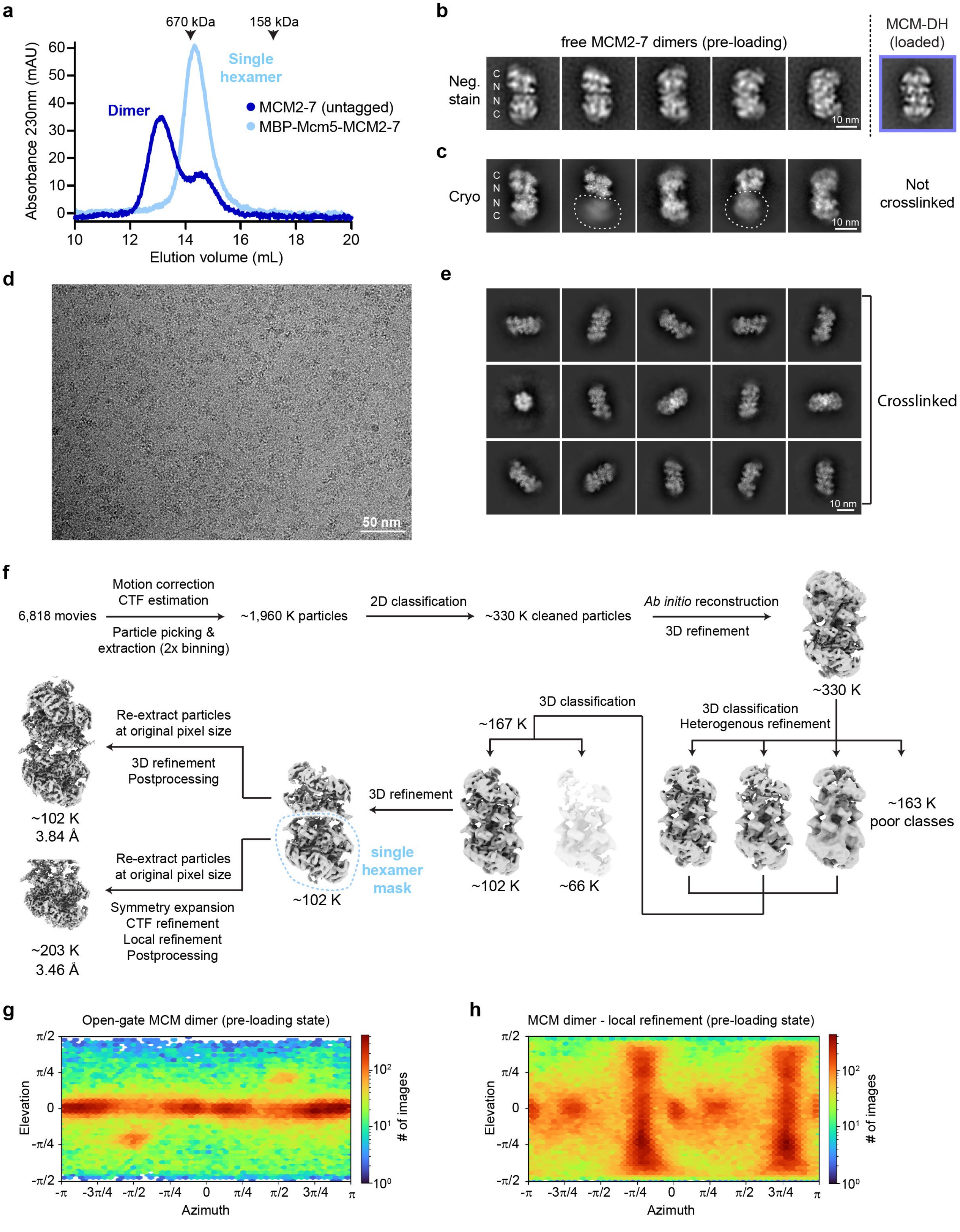
a
670 kDa
158 kDa
Absorbance 230nm (mAU)
Single hexamer
Dimer
MCM2-7 (untagged)
MBP-Mcm5-MCM2-7
Elution volume (mL)
b
free MCM2-7 dimers (pre-loading)
MCM-DH (loaded)
Neg. stain
10 nm
c
Cryo
Not crosslinked
d
50 nm
e
Crosslinked
f
6,818 movies
Motion correction
CTF estimation
Particle picking & extraction (2x binning)
~1,960 K particles
2D classification
~330 K cleaned particles
Ab initio reconstruction
3D refinement
~330 K
3D classification
Heterogenous refinement
~163 K poor classes
3D classification
~167 K
~102 K
~66 K
3D refinement
single hexamer mask
~102 K
Re-extract particles at original pixel size
3D refinement
Postprocessing
~102 K
3.84 Å
Re-extract particles at original pixel size
Symmetry expansion
CTF refinement
Local refinement
Postprocessing
~203 K
3.46 Å
g
Open-gate MCM dimer (pre-loading state)
Elevation
Azimuth
### of images
h
MCM dimer - local refinement (pre-loading state)
Elevation
Azimuth
### of images

**Extended Data Figure 7.** Open-ring human MCM2-7 hexamers can dimerize without loading onto DNA. **a**) Size exclusion chromatography of purified human MCM2-7 shows predominant dimerization of MCM2-7 hexamers (untagged). Fusing MBP to the N-terminus of Mcm5 prevents dimer formation. The elution of molecular weight markers (thyroglobulin – 670 kDa, $\gamma$-globulin – 158 kDa) is indicated by arrowheads. **b**) Negative-stain EM 2D class averages of MCM2-7 dimers, which are structurally distinct from loaded MCM double hexamers (class average shown for comparison). **c** to **h**) Cryo-EM data processing and validation of human MCM2-7 dimers. **c**) Cryo-EM 2D class averages of MCM2-7 dimers without crosslinking. Fuzzy density (outlined by dotted line) indicates conformational flexibility between both hexamers in the MCM dimer and prevented 3D reconstruction at high resolution. **d**) Cryo-EM image and **e**) 2D cryo-EM class averages of crosslinked (with glutaraldehyde) MCM dimers. **f**) Cryo-EM data processing workflow for 3D reconstruction. **g** and **h**) Angular distribution plots for MCM2-7 dimers (in **g**, C1 refined) and the locally refined, symmetry expanded MCM2-7 hexamer (in **h**). We note that although MCM2-7 was mixed with Cdt1 prior to cryo-EM sample preparation, no density is observed for this licensing factor, consistent with prior findings that the human proteins, unlike the *S. cerevisiae* counterparts[5,52], do not stably co-associate into an MCM2-7•Cdt1 heptamer[53].

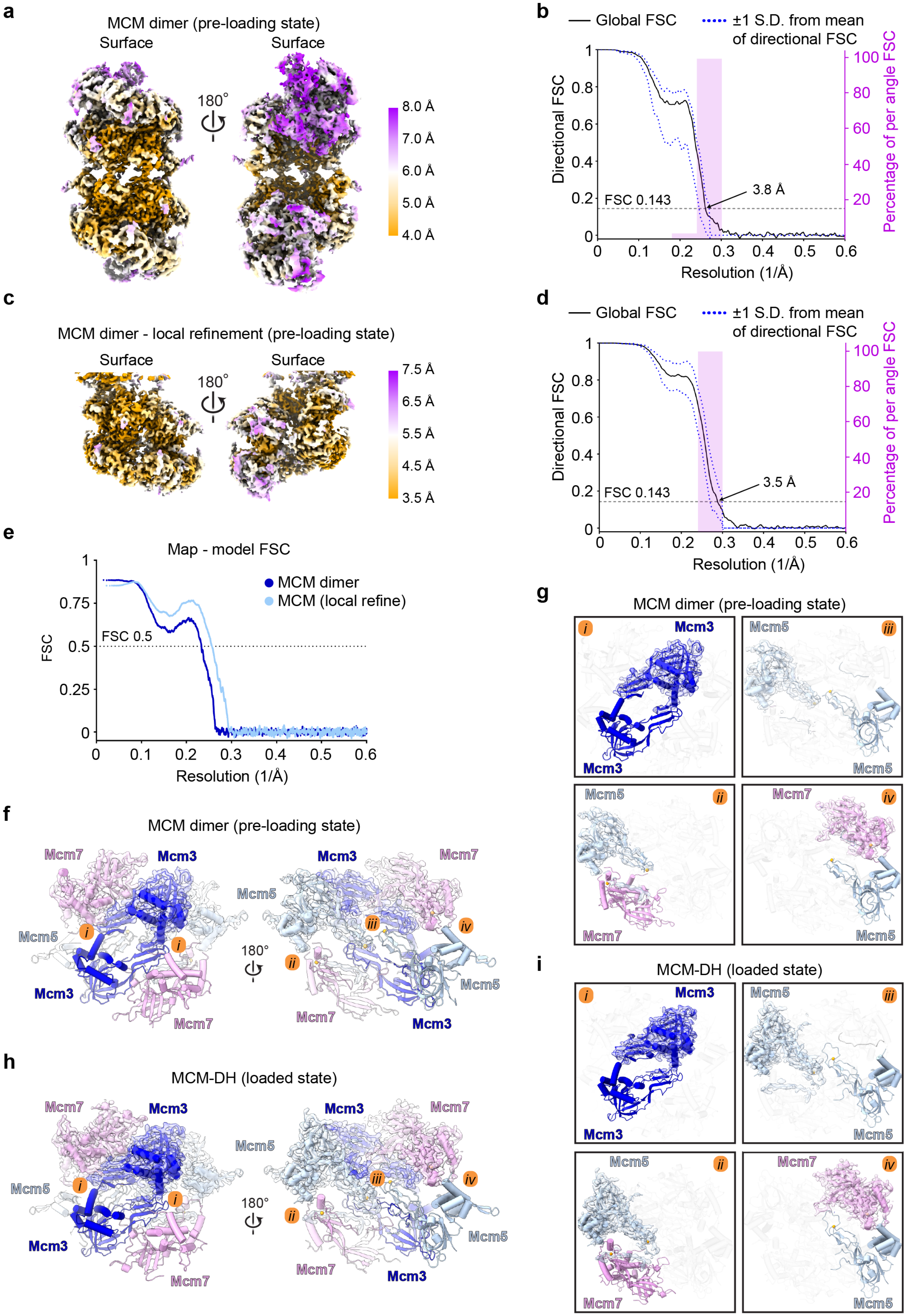

a
MCM dimer (pre-loading state)
Surface
Surface
180°
8.0 Å
7.0 Å
6.0 Å
5.0 Å
4.0 Å
b
Global FSC
±1 S.D. from mean of directional FSC
Directional FSC
Percentage of per angle FSC
FSC 0.143
3.8 Å
Resolution (1/Å)
c
MCM dimer - local refinement (pre-loading state)
Surface
Surface
180°
7.5 Å
6.5 Å
5.5 Å
4.5 Å
3.5 Å
d
Global FSC
±1 S.D. from mean of directional FSC
Directional FSC
Percentage of per angle FSC
FSC 0.143
3.5 Å
Resolution (1/Å)
e
Map - model FSC
MCM dimer
MCM (local refine)
FSC
FSC 0.5
Resolution (1/Å)
f
MCM dimer (pre-loading state)
Mcm7
Mcm3
Mcm5
Mcm7
180°
g
MCM dimer (pre-loading state)
Mcm3
Mcm5
Mcm7
h
MCM-DH (loaded state)
Mcm7
Mcm3
Mcm5
180°
i
MCM-DH (loaded state)
Mcm3
Mcm5
Mcm7

**Extended Data Figure 8.** Resolution estimation of MCM2-7 cryo-EM reconstructions in the pre-loading state and structural comparison with the loaded MCM double hexamer. **a** and **c**) Surface views of sharpened cryo-EM maps of full (in **a**) and locally refined (in **c**) MCM dimers. **b** and **d**) 3D Fourier shell correlation (FSC) plots for corresponding cryo-EM 3D reconstructions (full dimer map in **b** and locally refined map in **d**). **e**) FSC curves comparing cryo-EM maps and refined models. **f** to **i**) The MCM dimer is stabilized by similar interactions between Mcm3, Mcm5, and Mcm7 as in the loaded MCM double hexamer. **f** and **g**) Mcm5/3/7 dimerization interface in the open-gate MCM dimer. A structure overview is shown in **f** and zoomed views of the contact sites between subunits in the two hexamers (*i-iv*) in **g**. **h** and **i**) Mcm5/3/7 dimerization interface in the loaded MCM double hexamer. A structure overview is shown in **h** and zoomed views of the contact sites between subunits in the two hexamers (*i-iv*) in **i**. MCM subunits are shown as cartoon, with transparent cryo-EM map density overlaid in one of the hexamers in **f** to **i**.

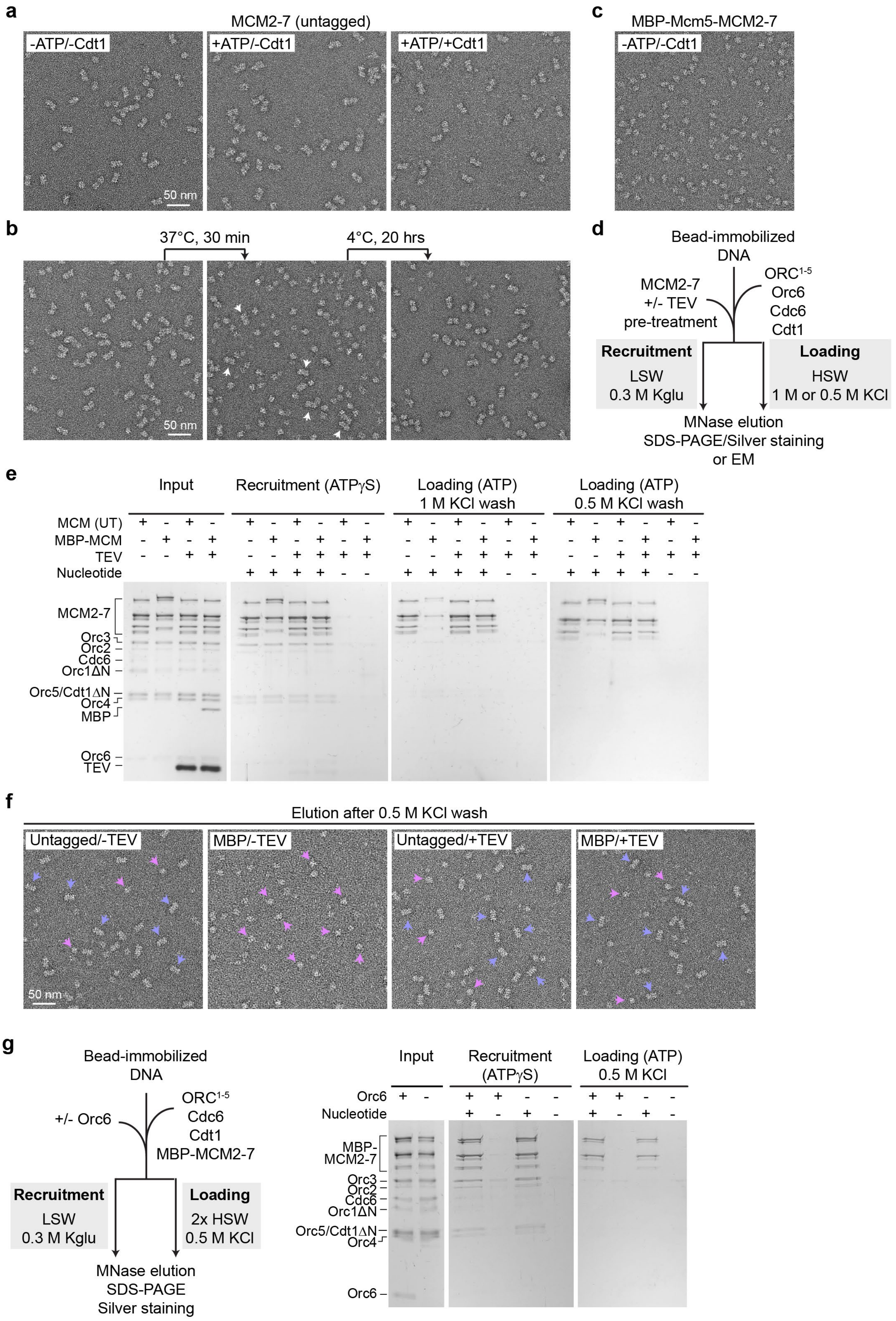
a
MCM2-7 (untagged)
-ATP/-Cdt1
+ATP/-Cdt1
+ATP/+Cdt1
50 nm
b
37°C, 30 min
4°C, 20 hrs
50 nm
c
MBP-Mcm5-MCM2-7
-ATP/-Cdt1
d
Bead-immobilized DNA
MCM2-7 +/- TEV pre-treatment
ORC1-5 Orc6 Cdc6 Cdt1
Recruitment
LSW 0.3 M Kglu
Loading
HSW 1 M or 0.5 M KCl
MNase elution SDS-PAGE/Silver staining or EM
e
Input
Recruitment (ATPγS)
Loading (ATP) 1 M KCl wash
Loading (ATP) 0.5 M KCl wash
MCM (UT)
MBP-MCM
TEV
Nucleotide
MCM2-7
Orc3
Orc2
Cdc6
Orc1ΔN
Orc5/Cdt1ΔN
Orc4
MBP
Orc6
TEV
f
Elution after 0.5 M KCl wash
Untagged/-TEV
MBP/-TEV
Untagged/+TEV
MBP/+TEV
50 nm
g
Bead-immobilized DNA
+/- Orc6
ORC1-5 Cdc6 Cdt1 MBP-MCM2-7
Recruitment
LSW 0.3 M Kglu
Loading
2x HSW 0.5 M KCl
MNase elution SDS-PAGE Silver staining
Input
Recruitment (ATPγS)
Loading (ATP) 0.5 M KCl
Orc6
Nucleotide
MBP-MCM2-7
Orc3
Orc2
Cdc6
Orc1ΔN
Orc5/Cdt1ΔN
Orc4
Orc6

**Extended Data Figure 9.** Human MCM2-7 can dimerize in the absence of other loading factors. **a**) Recombinant human MCM2-7 forms dimers irrespective of the presence of nucleotide or Cdt1. Incubations were done on ice. Subregions of negative-stain electron micrographs are shown. **b**) MCM2-7 dimers and monomers are in an equilibrium that is regulated by temperature. Incubation times and temperatures are indicated, starting from protein at 4ºC. While higher temperatures shift the equilibrium towards monomers, a subset of MCM2-7 remains in the dimeric form (arrowheads). Subregions of negative-stain electron micrographs are shown. **c**) Recombinant human MCM2-7 with an MBP-TEV-tag at the N-terminus of Mcm5 is exclusively monomeric (also on ice). A subregion of an electron micrograph of negatively stained MBP-Mcm5-MCM2-7 particles is shown. **d** to **g**) MBP at the N-terminus of Mcm5 supports the loading of MCM single hexamers onto DNA but not double hexamer formation. **d**) Summary of experimental workflow. **e**) MBP fused to the N-terminus of Mcm5 abrogates the loading of salt-resistant (1 M KCl) MCM complexes, but MBP removal by pre-treatment of MCM2-7 with TEV restores it. However, MBP-MCM is retained on DNA after washes with 0.5 M KCl buffer. Silver-stained SDS-PAGE gels of inputs and elutions from recruitment and loading reactions performed with MCM2-7 assemblies containing untagged Mcm5 or MBP-tagged Mcm5, either with or without pre-incubation of MCM2-7 with TEV, are shown. The residual MBP-MCM retained on DNA after 1 M salt wash are single MCM hexamers (see panel **f**). We note that several ORC subunits (Orc1ΔN, Orc2, Orc6) and Cdc6 tend to stain weakly with silver nitrate. **f**) Subregions of negative-stain electron micrographs of elutions from loading reactions after 0.5 M KCl wash. Only single MCM2-7 hexamers (magenta arrowheads) are seen with MBP-Mcm5, while MCM2-7 double hexamers are formed in other loading reactions (blue arrowheads). Kglu – potassium glutamate. **g**) Orc6 does not enhance loading of MCM2-7 single hexamers. Left: Summary of the experimental setup. Right: Silver-stained SDS-PAGE gels of inputs, recruitment, and loading reactions performed with MBP-MCM2-7 (MBP on Mcm5 N-terminus) in the absence or presence of Orc6. Note that +Orc6 lanes were also used for **Fig. 5g**.

**Extended Data Video 1.** Conformational flexibility in MO-like loading intermediates. Image sequence of MO-like 2D class averages (with the same MCM orientation) after alignment to the MCM density.

**Extended Data Table 1.** Summary of cryo-EM data collection, refinement, and validation statistics

| **Sample** | **Loaded MCM** (dataset 1) | **Loaded MCM** (dataset 2) | **Isolated MCM2-7** |
|---|---|---|---|
| **EM data collection and processing:** | | | |
| Microscope | Titan Krios | Titan Krios | Titan Krios |
| Camera | K3 | K3 | K3 |
| Voltage (kV) | 300 | 300 | 300 |
| Magnification | x105,000 | x105,000 | x105,000 |
| Frames (no.) | 40 | 40 | 40 |
| Total electron dose ($e^-/Å^2$) | 50.96 | 49.86 | 50.96 |
| Electron dose rate ($e^-/Å^2/s$) | 36.12 | 34.67 | 36.12 |
| Calibrated pixel size (Å) | 0.832 | 0.832 | 0.832 |
| Defocus range (μm) | -1.0 to -2.0 | -1.0 to -2.0 | -1.0 to -2.0 |
| Movies | 5,475 | 13,952 | 6,818 |
| Initial picks (no.) | 628,370 | 1,343,716 | 1,960,293 |

| | Loaded MCM-DH | Loaded MCM-SH | MCM dimer | Open-gate MCM (locally refined) |
|---|---|---|---|---|
| | PDB 8W0F<br>EMD-43708 | PDB 8W0E<br>EMD-43707 | PDB 8W0G<br>EMD-43709 | PDB 8W0I<br>EMD-43710 |
| Refined particles (no.) | 100,748 | 18,534 | 101,722 | 203,444 |
| Symmetry imposed | C1 | C1 | C1 | C1 (C2 symmetry expanded) |
| Global resolution (Å) | | | | |
| FSC 0.5 (masked) | 3.22 | 4.04 | 4.23 | 3.93 |
| FSC 0.143 (masked) | 2.75 | 3.35 | 3.84 | 3.49 |
| Map sharpening B factor ($Å^2$) | N/A (DeepEMhancer used) | N/A (DeepEMhancer used) | N/A (DeepEMhancer used) | N/A (DeepEMhancer used) |
| **Model refinement and validation:** | | | | |
| Initial model | 7W1Y | 7W1Y | 7W1Y | 7W1Y |
| Model composition | | | | |
| Non-hydrogen atoms | 63,321 | 30,082 | 56,920 | 28,381 |
| Protein residues | 7,721 | 3,647 | 7,134 | 3,557 |
| DNA residues | 94 | 50 | 0 | 0 |
| Ligands (ATP, ADP, Mg, Zn) | 2, 8, 10, 10 | 1, 4, 5, 5 | 8, 2, 10, 10 | 4, 1, 5, 5 |
| Root mean square deviation | | | | |
| Bond lengths (Å) | 0.003 | 0.002 | 0.003 | 0.003 |
| Bond angles (°) | 0.532 | 0.535 | 0.611 | 0.491 |
| B factors ($Å^2$) | | | | |
| Protein | 102.19 | 121.36 | 288.10 | 173.08 |
| DNA | 73.80 | 96.83 | N/A | N/A |
| Ligands | 92.22 | 104.53 | 249.25 | 157.70 |

| | | | | |
|---|---|---|---|---|
| Ramachandran plot | | | | |
| % favored | 95.02 | 94.98 | 94.96 | 95.63 |
| % allowed | 4.98 | 5.02 | 5.04 | 4.37 |
| % outliers | 0.00 | 0.00 | 0.00 | 0.00 |
| Rotamer outliers (%) | 1.08 | 0.63 | 0.51 | 0.03 |
| MolProbity | | | | |
| Clashscore | 6.31 | 6.90 | 5.46 | 4.48 |
| MolProbity score | 1.72 | 1.73 | 1.64 | 1.53 |
| Model-map comparison | | | | |
| $CC_{mask}$ | 0.80 | 0.80 | 0.71 | 0.80 |
| $FSC_{model/map}$ 0.5 | 3.2 | 3.8 | 4.4 | 3.9 |
